# Non-homologous end-joining is implicated in rapid and efficient caspofungin adaptation in *Nakaseomyces glabratus*

**DOI:** 10.64898/2026.11.03.763960

**Authors:** Laetitia Maroc, Nicholas C. Gervais, Meea Fogal, Eileen Bates, Garrett Westmacott, Peter McQueen, Stuart McCorrister, Christopher Grant, Sydney Johnson, Rebecca S. Shapiro

## Abstract

Antifungal resistance in *Nakaseomyces glabratus* is a major clinical challenge, yet the cellular processes that shape its evolution remain poorly understood. Here, we used experimental evolution to investigate the role of the error-prone double-strand break repair pathway, non-homologous end joining (NHEJ), in adaptation to fluconazole and caspofungin. Wild-type and NHEJ-deficient (*lig4Δ*) populations were evolved under antifungal selection for 25 days. Fluconazole resistance emerged rapidly in both genetic backgrounds and was associated with recurrent *PDR1* mutations and aneuploidies. In contrast, caspofungin adaptation was delayed and less efficient in *lig4Δ* populations, with several lineages failing to survive. All wild-type caspofungin-evolved isolates carried canonical *FKS2* hotspot mutations, whereas *lig4Δ* isolates predominantly acquired resistance through alternative genetic pathways. Caspofungin adaptation also frequently conferred cross-resistance to fluconazole and was accompanied by changes in the abundance of ergosterol-associated proteins. Together, these findings implicate NHEJ in promoting efficient evolution of FKS-mediated echinocandin resistance and demonstrate that DNA repair capacity can shape antifungal resistance trajectories in *N. glabratus*.

## Introduction

The annual incidence of invasive human fungal infections globally is estimated at approximately 6.5 million, with 2.6 million deaths directly attributable to these infections^1^. Among these, *Candida* species are major contributors to invasive mycoses, with *Candida*-associated bloodstream infections often linked to mortality rates exceeding 60%, though variation exists across locations^1^. In the absence of approved vaccines for human fungal diseases, clinicians rely heavily on prophylactic and post-infection antifungal therapeutics^2^. Unfortunately, the current arsenal of antifungal drugs for treating invasive *Candida* infections is extremely limited, and new antifungals have traditionally taken a long time to gain approval for human use^3^. In addition, the close evolutionary relationship between fungal and human cells has made it difficult to develop antifungal drugs that are both effective and non-toxic, while increasing rates of drug resistance amongst fungal pathogens, partly driven by the improper use of antifungals in agriculture and clinical settings, have further reduced the efficacy of existing treatments^4,5^. Although several new antifungal compounds are currently in clinical trials^6,7^, progress remains slow. Thus, there is an urgent need to elucidate the molecular mechanisms underlying antifungal drug resistance, understand how these resistance traits emerge, and identify which cellular processes contribute to their development.

Among *Candida* species, *Nakaseomyces glabratus* (formerly *Candida glabrata*) is particularly problematic due to its inherent low susceptibility to azoles, especially fluconazole, and the emergence of multidrug-resistant (MDR) isolates^8^. Unlike other *Candida* species, in which azole resistance is often acquired, *N. glabratus* exhibits intrinsically low susceptibility, rendering fluconazole frequently ineffective even at therapeutic concentrations^9,10^. At the molecular level, mechanisms driving decreased azole susceptibility in *N. glabratus* are diverse. This phenotype has mainly been attributed to gain-of-function mutations in the master transcriptional regulator *PDR1,* leading to constitutive overexpression of multidrug efflux pumps such as *CDR1,* and structural variations in the azole target enzyme Erg11^11–14^. In addition, genomic instability plays a crucial role in environmental adaptation and has been linked to antifungal resistance. For example, upon azole exposure, cells frequently acquire an extra copy of Chromosome E, which harbours *ERG11*, thereby increasing the dosage of the azole target enzyme^15,16^. Similarly, despite the echinocandin class of antifungals such as caspofungin being first-line treatment options, resistance readily emerges in *N. glabratus* and is often associated with mutations in hot-spot regions of *FKS1* and *FKS2,* which encode subunits of β-1,3-glucan synthase, therefore altering drug binding and reducing susceptibility^17–19^. Although polyene resistance remains rare, amphotericin B toxicity limits its clinical use, leaving few viable treatment options^20^. The growing prevalence of MDR *N. glabratus* isolates, resistant to both azoles and echinocandins, poses a major clinical challenge and underscores the urgent need to better understand the mechanisms that drive antifungal drug resistance in this species.

Beyond these well-characterized mechanisms, there is growing evidence that DNA damage repair pathways influence antifungal drug susceptibility in both *Candida albicans* and *N. glabratus*^21–23^. Notably, deletion of genes involved in double-strand break (DSB) repair and mismatch repair in *C. albicans* appeared to alter genomic stability and susceptibility to fluconazole, though the specific roles of different repair pathways remain unclear^21^. A recent study showed that the putative error-prone polymerase, Rev1, mediates DNA damage and is associated with caspofungin resistance in *C. albicans* ^23^. While orthologous mismatch repair factors have been implicated in *N. glabratus*, the role of DSB repair pathways, especially non-homologous end joining (NHEJ), in antifungal drug resistance remains underexplored^22^. Thus, while many mutations have been identified as either directly contributing to or associated with drug-response phenotypes in *N. glabratus*, the exact mechanisms underlying their emergence, including the potential involvement of DSB repair pathways, remain poorly understood^16^.

NHEJ repairs double-strand breaks independently of a homologous template and, as a consequence, frequently introduces small insertions or deletions (INDELs) at the repaired locus. In short, NHEJ depends on a small set of core proteins that rapidly bind and repair broken DNA ends ^24^. The Ku70-Ku80 heterodimer recognizes and protects DSB ends, while accessory factors such as Nej1 and Pol4 help align the ends using microhomology and process the DNA for repair. Dnl4 ligase, in complex with Lif1 and Nej1, performs the final ligation. Together, these proteins enable fast but error-prone DSB repair. In *N. glabratus*, NHEJ is highly efficient compared with other yeasts^25,26^, and key genes in this pathway, such as *DNL4* (also called *LIG4*, *CAGL0E02695g*), show evidence of positive selection ^27^, suggesting an important evolutionary role in DNA repair and adaptation. Given that antifungal exposure induces stress conditions that can cause DNA damage^28^, understanding how NHEJ contributes to genome plasticity and drug adaptation may yield novel insights into the emergence of antifungal resistance in this species.

We therefore used adaptation laboratory evolution (ALE) to investigate the role of NHEJ in *N. glabratus’s response* to two of the most commonly used antifungal drugs in clinical treatment: fluconazole and caspofungin. ALE has represented a powerful strategy for studying the dynamics and mechanisms of antifungal drug resistance in *N. glabratus* and many other fungal pathogens ^13,16,29,30^. Specifically, we expose wild-type (WT) and NHEJ-deficient (Δlig4) populations of *N. glabratus* to fluconazole and caspofungin over 25 days. We demonstrate a putative role for NHEJ in potentiating caspofungin resistance in *N. glabratus*. We combine whole-genome sequencing (WGS) and proteomics to identify aneuploidies, single-nucleotide polymorphisms (SNPs), and differentially abundant proteins associated with drug-resistant populations, and capture adaptation strategies at both the single-colony and population levels. We also monitor the stability of drug-susceptibility phenotypes in our evolved populations by passaging them under neutral conditions. We find that many populations evolved in the presence of caspofungin demonstrate cross-adaptation to fluconazole, though this cross-adaptation appears less stable than resistance to caspofungin. Finally, we characterize the involvement of acquired resistance mutations in fungal virulence and broader stress responses, showing that adaptation to antifungal drugs often leads to trade-offs. Taken together, our results highlight the molecular basis of diverse resistance mechanisms from a multi-omics perspective and comprehensively profile properties of the resulting mutant populations relevant to human fungal infection and clearance.

## Results

### NHEJ is involved in rapid and efficient caspofungin adaptation in N. glabratus

We first explored the role of the DNA repair pathway, NHEJ, in antifungal drug adaptation in *N. glabratus.* For this, we initiated an adaptive laboratory experiment (ALE) in which a wild-type strain (WT) and a NHEJ-deficient mutant strain (*lig4*Δ) were passaged in triplicate populations and exposed to increasing concentrations of the antifungal fluconazole (FLZ), or a constant concentration of the antifungal caspofungin (CASP) for 25 days with daily passaging (Fig.1).

**Figure 1.**
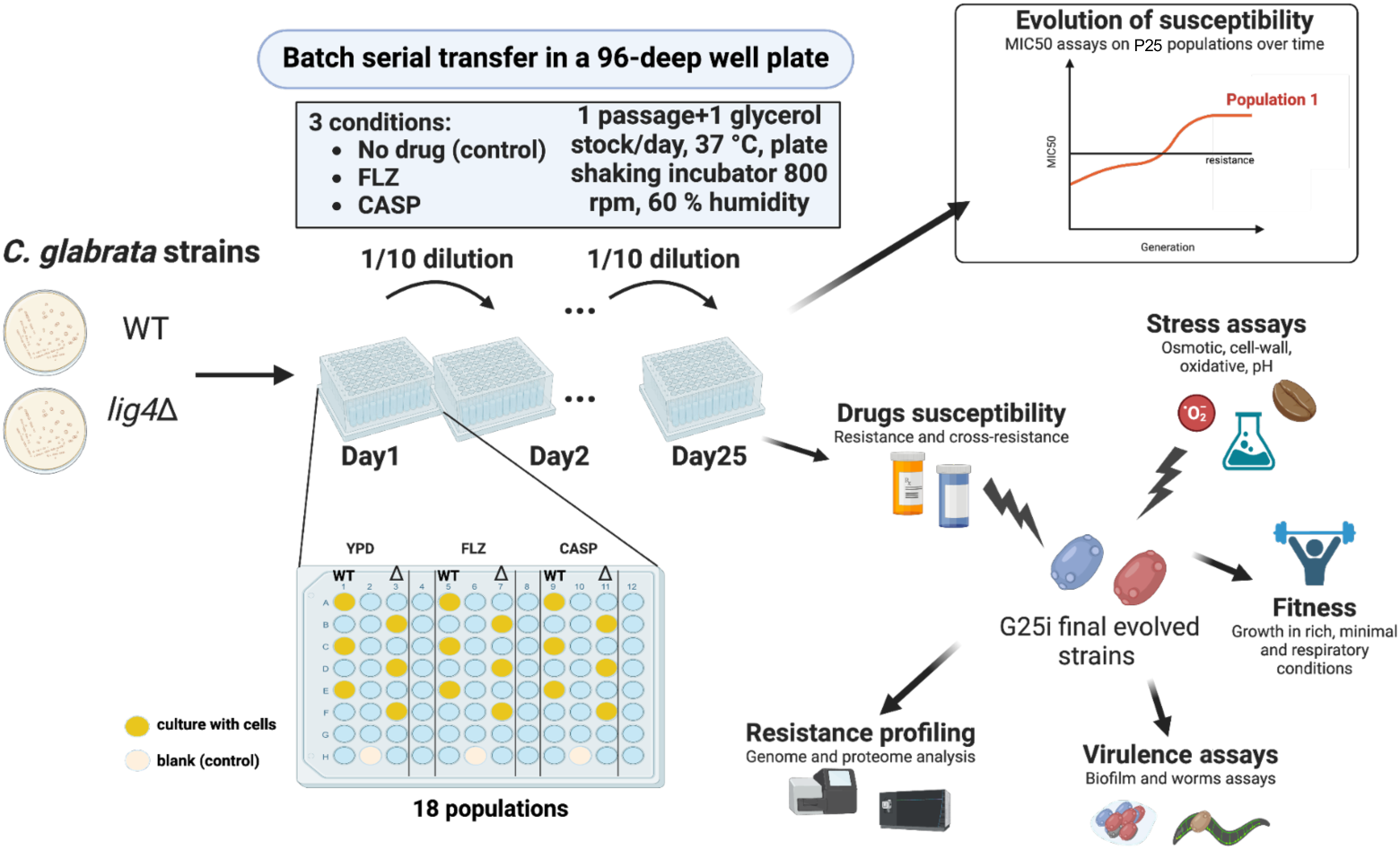
Experimental design of the adaptive laboratory experiment (ALE). A wild-type (WT) strain and an NHEJ-deficient mutant strain (*lig4Δ*, Δ) were subjected to drug treatments (CASP, caspofungin; FLZ, fluconazole) or grown in YPD as a control for 25 days with daily passaging. Before each passage, a glycerol stock was prepared from each population. Drug susceptibility of the evolved populations was assessed throughout the experiment. At the end of the experiment, a single colony from each population was isolated for further strain characterization.

During the experiment, we observed that adaptation to FLZ remained stable after passage 9 (P9), and at P15, we increased the FLZ concentration from 160 µg/mL to 640 µg/mL (equivalent to twice the MIC50 of most FLZ-evolved populations at P15) (Fig. 2A). In contrast, CASP-evolved populations exhibited a continuous increase in drug adaptation over time, despite the CASP concentration remaining constant throughout the experiment (Fig. 2B). As a control, both WT and *lig4Δ* strains were passaged in triplicate populations in rich media without antifungal agents (referred to as neutrally-evolved populations). In total across all passages, our experiment yielded 150 neutrally-evolved and 300 drug-evolved population samples.

**Figure 2.**
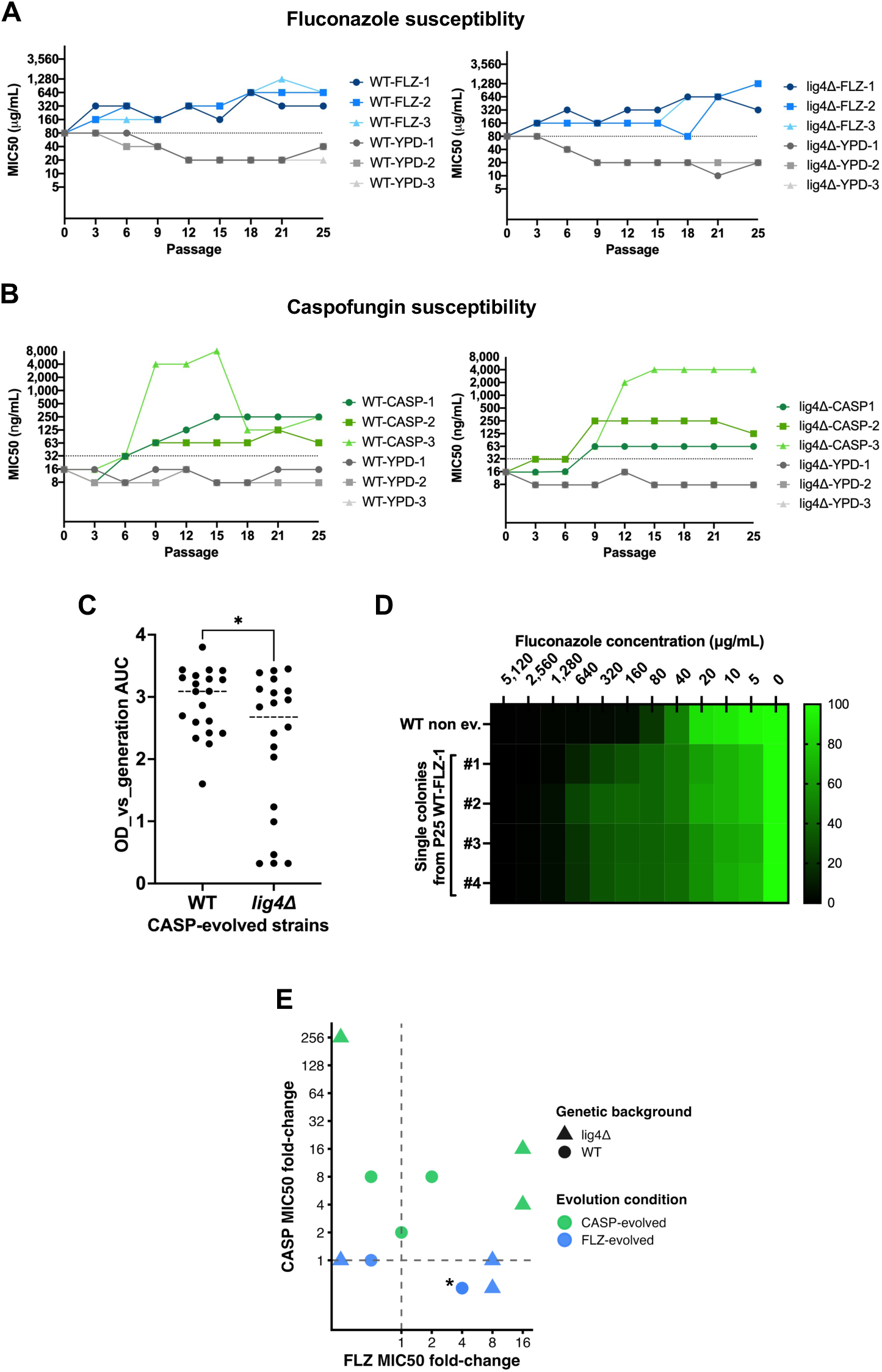
Caspofungin and fluconazole susceptibility of drug-evolved strains. **(A, B)** Drug susceptibility profiles of YPD- and FLZ-evolved strains to fluconazole (FLZ) **(A)** and of YPD- and CASP-evolved strains to caspofungin (CASP) **(B).** Susceptibility was assessed every three passages and at P25 for each evolved population recovered from glycerol stocks. The dotted lines represent the MIC50 of the ancestral WT and *lig4Δ* strains. **(C)** Fitness of the CASP-evolved populations in the presence of CASP. The OD600 of 20 CASP-evolved populations per strain was measured every 24 hours for 6 days (six passages). The graph represents the area under the curve (AUC) of OD600 over time. Each dot represents an individual population, and the bar indicates the median. *: Pvalue = 0.02 (unpaired, two-tailed, Welch’s t-test). **(D)** Fluconazole MIC50 assay of the ancestral WT non-evolved strain and single colonies isolated from the P25 WT-FLZ-1 population. Strains were grown at 37 °C in YPD containing a gradient of fluconazole concentrations, and OD600 was measured after 24 h of growth. **(E)** MIC50 fold changes of P25-sg drug-evolved strains relative to the ancestral strain. The vertical and horizontal dashed lines indicate the resistance thresholds for fluconazole and caspofungin, respectively. The blue data point marked with an asterisk represents two independent strains that share identical fluconazole and caspofungin MIC50 fold-change values and therefore overlap completely on the plot.

We sought to determine how rapidly wild-type and NHEJ-deficient strains could adapt to antifungal treatment and to evaluate the dynamics of this adaptation over the course of the experiment. We thus assessed the antifungal drug susceptibility of these strain lineages over time by performing MIC assays on all of the neutrally-evolved and drug-evolved populations from every third passage (Fig. 2A and Fig. 2B; Table S1 and Table S2). We define a ‘drug-adapted’ population as an evolved population with a MIC50 at least 2-fold higher than that of its ancestor strain. Since the non-evolved WT and *lig4*Δ ancestor strains have identical MIC50 values for both FLZ and CASP (Table S3), this enables direct comparison of drug susceptibility across all populations throughout the experiment.

Over the course of passaging in the absence of drugs, susceptibility to FLZ and CASP increased slightly but steadily, with MIC50 values decreasing by 2-fold for FLZ and 1-fold for CASP relative to the ancestral strains in both WT and *lig4Δ* populations (Fig.2A and Fig. 2B). As expected, none of the neutrally evolved populations ever developed resistance to the corresponding antifungal treatment, confirming that passaging alone does not lead to antifungal adaptation. For the FLZ-evolved populations, we found that both WT and *lig4*Δ populations adapted to FLZ treatment in as few as 3 days (Fig. 2A), confirming *N. glabratus*’s ability to rapidly adapt to growth in the presence of azole drugs. When the FLZ concentration was increased at P15 from 160 µg/mL to 640 µg/mL, all populations were able to adapt to this higher drug concentration by P21 (Fig. 2A).

Wild-type strains exposed to CASP developed resistance by P6, while two out of three *lig4Δ* populations did not develop resistance until P9. This indicated that *N. glabratus* adapted more slowly to CASP than to FLZ, with the delay possibly being more pronounced in the *lig4Δ* mutant. Nevertheless, this trend should be interpreted cautiously, as only three independent populations were analyzed for each genetic background. Although drug concentrations remained constant throughout the experiment (or until P15 for FLZ treatment), most populations adapted to drug levels higher than those used during passaging. Notably, CASP treatment resulted in a diverse range of resistance profiles among adapted strains, with MIC50 values ranging from 64 ng/mL to 4,000 ng/mL (Fig. 2B and Table S2).

To achieve sufficient statistical power to formally compare adaptation rates between CASP-evolved *lig4Δ* and WT populations, we increased the number of independently evolving lineages to 20 per genetic background. This was achieved by performing an additional evolution experiment with 17 biological replicates of each strain in YPD containing 32 ng/mL CASP for 6 days and combining these data with those from the initial experiment. We monitored their fitness in CASP over time by measuring the OD600 of each population before each passage (Table S4). We then calculated the area under the OD600-versus-passage curve (OD_vs_Passage AUC). As shown in Fig. 2C, WT populations exhibited significantly higher fitness in CASP than *lig4Δ* populations (Welch’s *t*-test, p-value = 0.02). Additionally, four *lig4Δ* populations did not survive the experiment (populations 5, 15, 18, and 19, Table S4), whereas all WT populations persisted. These results suggest that *lig4Δ* populations have a reduced capacity for efficient adaptation to CASP, indicating that while NHEJ is not essential, it may play an important role in CASP adaptation.

### Adaptation generates heterogeneous lineage-specific responses, whereas cross-adaptation is less stable

Next, we isolated single colonies (P25-sg strains) from each P25 population and determined their MIC50 values (Tables S5 and S6). Several evolved populations were heterogeneous with respect to drug susceptibility, as susceptible colonies could still be recovered after 25 days of antifungal exposure from populations that displayed an overall resistant phenotype. This heterogeneity was particularly evident in two fluconazole-evolved populations, WT-FLZ-1 and lig4Δ-FLZ-1. Although both P25 populations were classified as resistant (MIC50 = 320 µg/mL), we were unable to isolate individual colonies with a similarly resistant phenotype. Instead, the recovered colonies had MIC50 values equal to or lower than those of their respective ancestral strains (ancestral MIC50 = 80 µg/mL; Table S5; Fig. 2D). Despite their reduced resistance, these isolates exhibited increased FLZ tolerance, displaying slow growth at inhibitory drug concentrations and reaching approximately 20% growth at 640 µg/mL FLZ, compared with ∼1% growth for the ancestral strains (Table S5; Fig. 2D).

These findings suggest that the WT-FLZ-1 and lig4Δ-FLZ-1 populations may contain a small subpopulation of highly resistant cells that were not recovered during single-colony isolation, and/or a larger population of tolerant cells whose collective growth contributes to the resistant phenotype observed at the population level ^31^. More broadly, these results suggest that adaptation to fluconazole can produce distinct resistance and tolerance phenotypes within the same evolved population. For subsequent experiments, we selected colonies with MIC50 values within one two-fold dilution of their corresponding P25 population, where possible, to ensure that individual isolates were representative of the population-level susceptibility phenotype (Tables S5 and S6). Although WT-FLZ-1-sg and lig4Δ-FLZ-1-sg did not meet this criterion, we retained these isolates for follow-up analyses because their stable tolerance phenotypes provided an opportunity to investigate an alternative mode of fluconazole adaptation.

We next evaluated cross-adaptation of the P25-sg strains to an antifungal drug not used during experimental evolution. Three of six CASP-resistant strains (WT-CASP-1-sg, lig4Δ-CASP-2-sg, and lig4Δ-CASP-3-sg) exhibited cross-resistance to FLZ, whereas none of the FLZ-evolved strains showed cross-resistance to CASP (Fig. 2E; Table S7). This pattern is consistent with previous reports of cross-resistance in laboratory-evolved *N. glabratus* strains ^16^. To determine the stability of these drug-response phenotypes, we passaged all drug-evolved P25-sg strains for six days in rich medium without antifungal selection, with daily transfers, and then reassessed susceptibility (Table S7). Resistance to the drug used during experimental evolution remained stable: all resistant strains retained MIC50 values within one-to two-fold dilution of their pre-passaging values. FLZ tolerance was also maintained in WT-FLZ-1-sg and lig4Δ-FLZ-1-sg. In contrast, cross-resistance was not always stable. WT-CASP-1-sg lost its FLZ cross-resistance after drug-free passaging, with its FLZ MIC50 decreasing eightfold, from 160 to 20 µg/mL.

### Evolving resistance to fluconazole leads to a larger fitness cost than caspofungin

During infection and persistence in a host, *N. glabratus* experiences variations in temperature, nutrient and oxygen availability, and pH, which challenge its survival^32^. Adaptation to environmental stressors is therefore key to survival and host invasion. However, adaptation to stressful growth conditions, like exposure to antifungal agents, can lead to fitness trade-offs ^16,33–35^. We therefore examined how drug-evolved strains responded to diverse stress conditions by exposing P25-sg lineages to multiple environmental challenges and assessing their fitness and stress resistance using serial-dilution spotting assays and growth-curve analyses (Fig. 3 and Fig. S1).

**Figure 3.**
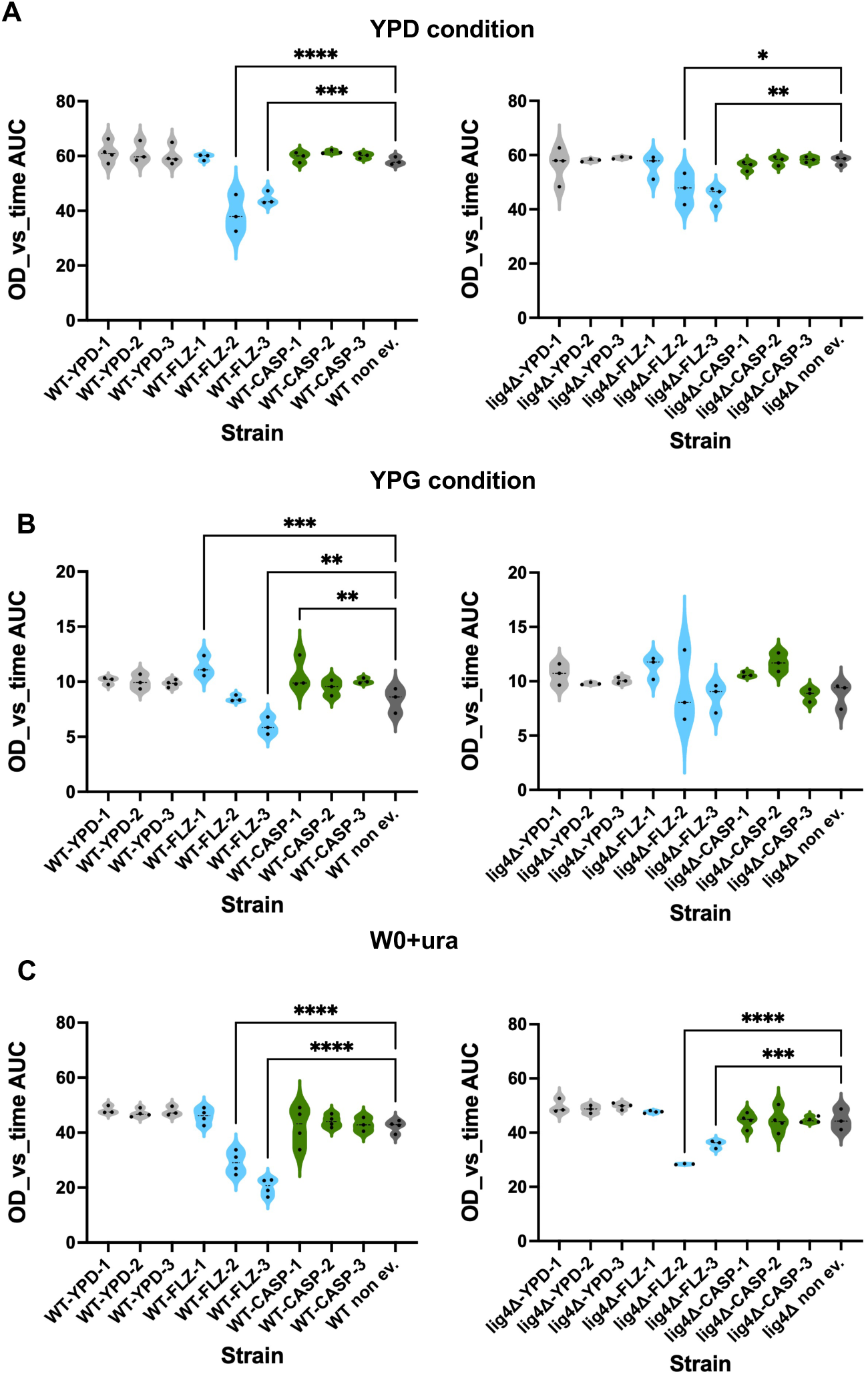
Impact of drug resistance on the fitness of P25-sg strains. Area under the curve (AUC) values from kinetic growth assays (OD600 over time for 24 hours). Fitness was assessed by measuring growth under rich (YPD) **(A)**, respiratory (YPG) **(B)**, and nutrient-limited (WO + ura) **(C)** conditions for 24 hours at 37 °C. Each dot represents one biological replicate, and the bar indicates the median. *P < 0.05, **P < 0.01, ***P < 0.0005, ****P < 0.0001 (ordinary one-way ANOVA followed by Dunnett’s multiple comparisons test using the corresponding non-evolved parental strain as the control: WT non-evolved for WT-evolved strains and *lig4Δ* non-evolved for *lig4Δ*-evolved strains).

P25-sg lineages were first tested for fitness in three different nutrient and carbon source conditions: rich medium (YPD), respiratory condition medium (YPG, containing glycerol rather than glucose as the carbon source, thereby favouring respiratory metabolism), and nutrient-limited medium (W0+ura) (Fig. 3). Neutrally evolved strains exhibited fitness comparable to ancestral non-evolved strains across all tested conditions, confirming that passaging in the absence of antifungal agents did not impact overall fitness. In contrast, all FLZ-adapted strains (both WT and *lig4Δ*) showed significant growth defects in non-respiratory conditions YPG (Fig. 3B), YPD, and W0+ura. However, except for WT-FLZ-3-sg, we observed no growth defects in YPG. Notably, two evolved strains, WT-FLZ-1-sg (tolerant to FLZ) and WT-CASP-1-sg (resistant to CASP), demonstrated enhanced fitness in YPG (Fig. 3B). This suggests that FLZ treatment may impair the ability to utilize glucose as a sole carbon source and that antifungal drug treatments, at least FLZ and CASP, can lead to improved fitness in respiratory growth conditions. We observed no fitness defects in CASP-evolved strains across all tested conditions (YPD, W0+ura, and YPG; Fig. 3).

We next tested the P25-sg strains for resistance to cell wall stressors (calcofluor white [CW] and caffeine), the osmotic stressor NaCl, the oxidative stressor hydrogen peroxide (H₂O₂), and extreme pH levels (acidic pH 3 and 5, and basic pH 10) (Fig. S1). Consistent with the fitness assay results, neutrally evolved strains exhibited stress sensitivity comparable to that of the ancestral non-evolved strains across all tested conditions (Fig. S1). All drug-adapted strains, except one CASP-evolved strain (WT-CASP-2-sg), showed increased sensitivity to at least one stress condition (Fig. S1). Stress sensitivity did not differ between WT and *lig4Δ* drug-evolved strains. However, only FLZ-resistant strains exhibited heightened sensitivity to NaCl, consistent with previous reports of NaCl sensitivity in FLZ-evolved *N. glabratus* strains^16^. Interestingly, WT-CASP-2-sg was the only strain that did not show increased susceptibility to any of the tested stress conditions; it even showed reduced sensitivity to 10 mM caffeine, a known cell wall stressor (Fig. S1). These findings suggest that while growth defects are frequently associated with drug resistance^16,33–35^, beneficial growth effects can also arise in specific cases.

### Virulence in a host infection model does not correlate with biofilm formation capacities

*N. glabratus* can adhere to various medical devices and human tissues via the formation of adherent biofilms^36^. Biofilm formation often plays a crucial role in *Candida* species’ susceptibility to antifungal agents and in host interactions by acting as a barrier that hinders fungal clearance^37^. Indeed, many studies have reported a link between *N. glabratus* biofilm formation and antifungal drug resistance^36,38,39^. We therefore sought to assess the biofilm capacity of P25-sg-evolved strains (both neutrally evolved and drug-evolved). We found that half of the drug-evolved P25-sg strains had significantly different biofilm-forming capacities than their respective ancestral non-evolved strains (Fig. 4A), suggesting that adaptation to antifungal drugs can coincide with changes in biofilm-forming ability in *N. glabratus*, though not consistently.

**Figure 4.**
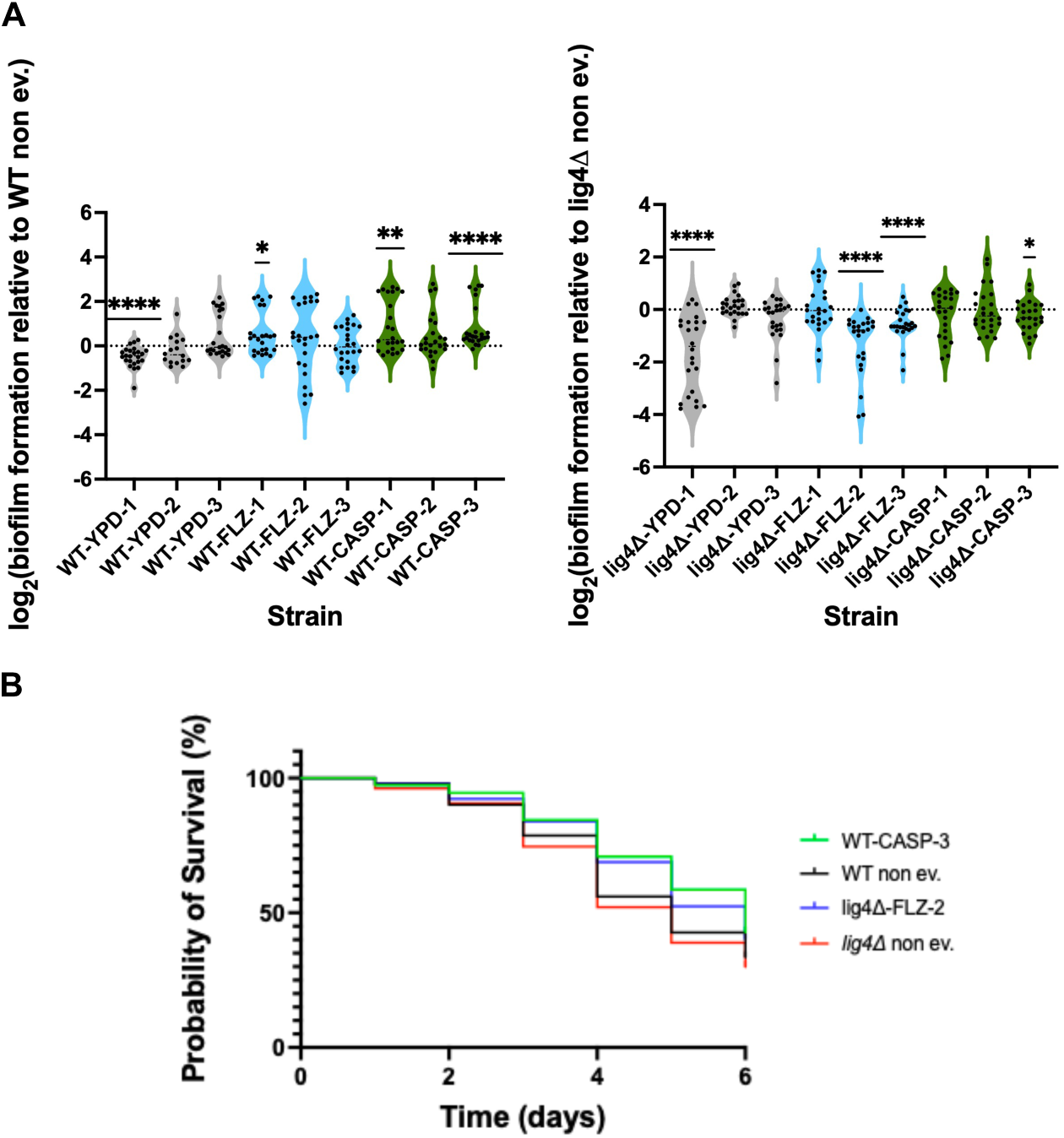
Impact of drug resistance on biofilm formation. **(A)** and pathogenicity **(B)** of P25-sg strains. **(A)** Biofilm formation was assessed using the tetrazolium salt (XTT) reduction assay in YPD. The relative biofilm formation was calculated as follows: (OD_490nm_/OD_600nm_ planktonic cells)_condition_/(OD_490nm_/OD_600nm_ planktonic cells)_ancestor strain_ where the condition strain is a P25-sg strain and the ancestor strain is the corresponding non-evolved parental strain. Dots represent individual biological replicates from three independent experiments, each containing 8 technical replicates (n=24 total apart from WT-YPD-2-sg where n=16), and the dotted line indicates a value of 0, corresponding to no change in biofilm formation relative to the ancestor. *P < 0.05, **P < 0.01, ***P < 0.0005, ****P < 0.0001 (Wilcoxon signed-rank test, compared with the hypothetical value of 0). **(B)** Pathogenicity of P25-sg strains assessed using a *Caenorhabditis elegans* infection model. Two drug-evolved P25-sg strains were evaluated: WT-CASP-3-sg, which exhibited increased biofilm formation, and lig4Δ-FLZ-2-sg, which exhibited reduced biofilm formation. Data are shown from three independent experiments.

Interestingly, the evolution of biofilm capacities differed markedly depending on the strains’ genetic background (WT or *lig4Δ*) but not on the specific prior drug exposure (FLZ or CASP). Compared to their ancestral non-evolved strains, WT-FLZ-1-sg, WT-CASP-1-sg, and WT-CASP-3-sg formed more robust biofilms, whereas lig4Δ-FLZ-2-sg, lig4Δ-FLZ-3-sg, and lig4Δ-CASP-3-sg exhibited reduced biofilm-forming capacity. The reduced biofilm formation in the *lig4Δ* drug-evolved strains may be a consequence of passaging rather than drug treatment, as two neutrally-evolved strains, WT-YPD-1-sg and lig4Δ-YPD-1-sg, also formed less biofilm than their ancestral non-evolved strain. However, none of the neutrally evolved strains developed a greater capacity to form biofilm after passaging, and WT and *lig4Δ* ancestral non-evolved strains exhibit identical biofilm-forming capacities (Fig. S2). It is thus likely that drug treatments contributed to increased biofilm-forming capacity, specifically in WT-derived strains.

Since biofilm formation can contribute to *Candida* species’ ability to evade the host immune system and thus their virulence, we next sought to determine whether variable biofilm-forming capacities of our drug-evolved isolates were associated with differences in virulence in a host model. Therefore, we exploited *Caenorhabditis elegans* as a model host for fungal infection^40,41^ and performed infection assays with two P25-sg strains: lig4Δ-FLZ-2-sg, which has a reduced capacity to form biofilm, and WT-CASP-3-sg, which forms more robust biofilms. Both strains had significantly reduced virulence in worms (Fig. 4B) compared to their ancestral, non-evolved strains (Log-rank Mantel-Cox test, p-value = 0.002 for WT strains and p-value = 0.0005 for *lig4Δ* strains). No significant differences were found between the WT and the *lig4Δ* non-evolved parental strains (Log-rank Mantel-Cox test, p-value = 0.26), suggesting that deletion of *LIG4* does not affect virulence in worms.

Together, this suggests that the mutations selected under antifungal exposure in the WT-evolved strains may differ from those selected in the *lig4Δ* strains, conferring distinct biofilm-formation capacities. Furthermore, acquired drug adaptations can reduce virulence, at least in a host infection model. It also suggests that, under the conditions tested, biofilm growth is uncoupled from virulence in *N. glabratus.* This decoupling of biofilm growth and virulence has been reported in *C. albicans*^41–43^, but to our knowledge, it has not been observed in *N. glabratus*.

### Fluconazole adaptation is associated with canonical resistance mutations and aneuploidies

We next profiled the resistance of P25-sg strains using a combination of targeted and whole-genome sequencing. Since FLZ resistance can arise through a variety of mutations and chromosomal rearrangements in *N. glabratus*^20^, we performed whole-genome sequencing (WGS) on the P25-sg FLZ-evolved strains to assess SNPs and INDELs (insertions/deletions) in genome coding regions, and aneuploidies. For FLZ-evolved strains, we thus sequenced a total of six strains (four FLZ-adapted strains: WT-FLZ-2-sg, WT-FLZ-3-sg, lig4Δ-FLZ-2-sg, and lig4Δ-FLZ-3-sg; and two FLZ-tolerant strains: WT-FLZ-1-sg and lig4Δ-FLZ-1-sg). As expected^16,44,45^, aneuploidies were common in FLZ-resistant strains, with at least one chromosome duplication common (chromosome C, E and F for WT-FLZ-2-sg and lig4Δ-FLZ-2-sg, and chromosome E for WT-FLZ-3-sg and lig4Δ-FLZ-3-sg; Fig. 5). The amplification of genes involved in drug susceptibility often underlies aneuploidy-mediated drug resistance in *Candida* species^46,47^. Indeed, many genes involved or linked to azole resistance in *N. glabratus* are encoded on these chromosomes. For instance, chromosome C harbours genes *HSP31* and *UPC2A*^14,48–50^ for which gain-of-function has previously been linked to FLZ resistance in *N. glabratus*. Chromosome F encodes *CDR2* and *ERG3,* while chromosome E encodes *ERG11*(fluconazole drug target), *TPO1*, and *ADA3*, genes that have been implicated in fluconazole susceptibility and resistance in *N. glabratus*^51^. Furthermore, chromosome F has been identified as a common aneuploidy associated with FLZ resistance in *N. glabratus*^16^.

**Figure 5.**
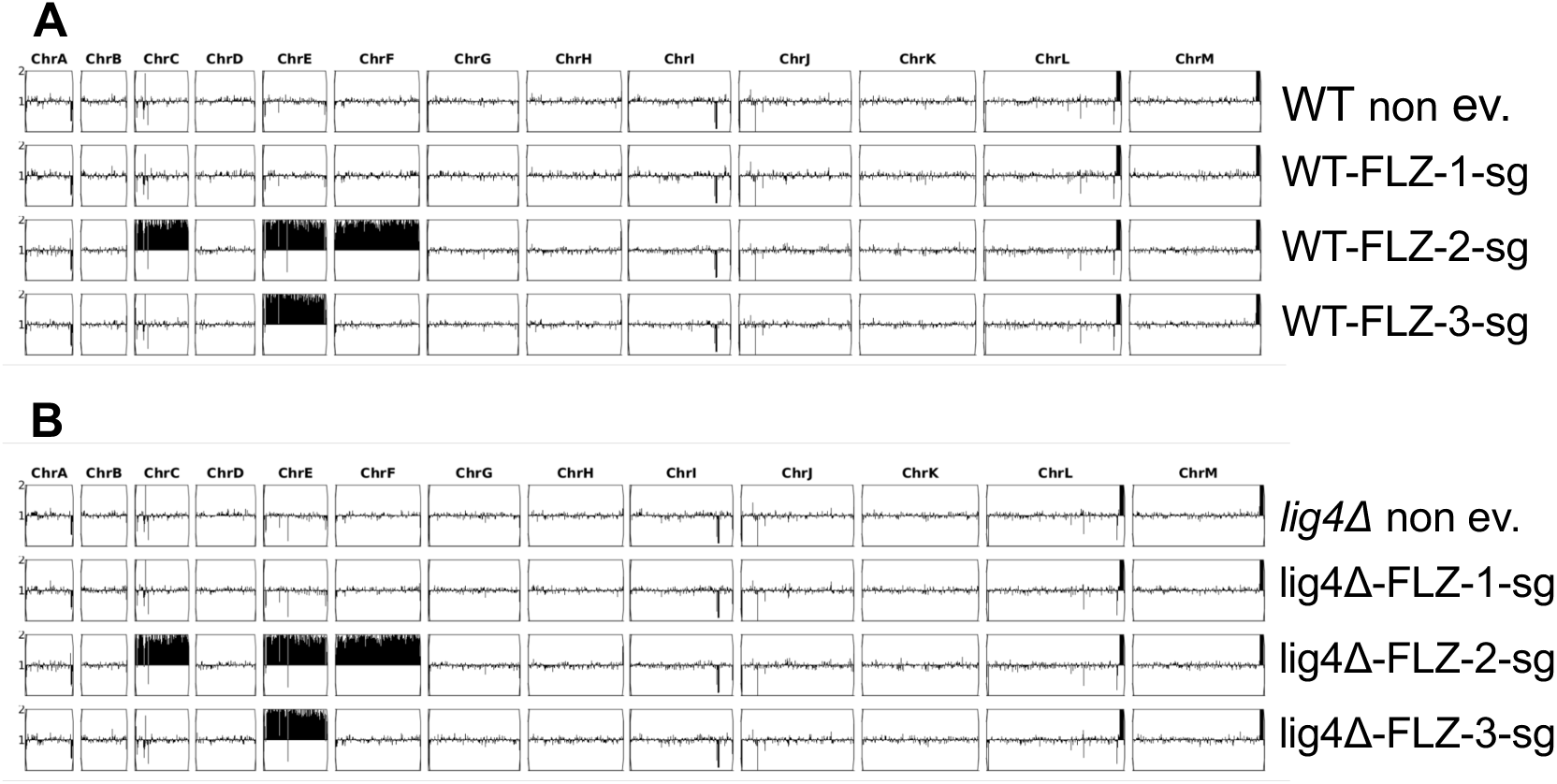
Ploidy status of FLZ-evolved strains and their corresponding non-evolved ancestral strains. For each genetic background (WT **(A)** and lig4Δ **(B)**), chromosome copy number was determined using the Ymap analysis tool. Black rectangles indicate whole-chromosome duplications.

In addition to the identified aneuploidies, 3 of 4 FLZ-adapted strains (WT-FLZ-2-sg, lig4Δ-FLZ-2-sg, and lig4Δ-FLZ-3-sg) carried a non-synonymous mutation in *PDR1*, which encodes the central transcriptional regulator of fluconazole susceptibility in *N. glabratus* (Document S1) and is common in fluconazole-resistant *N. glabratus* clinical isolates^19^. Interestingly, the only FLZ-adapted strain that did not contain a mutation in *PDR1*, WT-FLZ-3-sg, also did not contain mutations in any other well-known resistance genes, yet this strain has a similar resistance profile to those with a mutated *PDR1* (Document S1 and Fig. 5). However, WT-FLZ-3-sg is aneuploid for chromosome E and carries a few mutations in genes not previously associated with FLZ resistance, suggesting that chromosome E aneuploidy or additional mutations in genes not yet linked to fluconazole resistance may lead to adaptation to high levels of FLZ.

In the two FLZ-tolerant strains, WT-FLZ-1-sg and lig4Δ-FLZ-1-sg, we did not detect any large genomic rearrangements, *PDR1* mutations, or mutations in any established fluconazole resistance genes (Document S1 and Fig. 5). However, since we previously showed that tolerance was stable in these strains, the tolerance might be due to one or a combination of the identified mutations. WT-FLZ-1-sg contained a combination of mutated genes (*IZH3*, *PBL3,* and *FET3*) associated with antifungal resistance, fitness, or biofilm formation in *Candida* species (Table 1). Strain lig4Δ-FLZ-1-sg contains a higher number of mutated genes than other FLZ-evolved strains (Document S1, 11 genes vs. up to 5 genes in other strains), including one in *MSH2,* which is involved in DNA mismatch repair and whose deletion leads to an increased frequency of drug-resistant mutants ^22^. Altogether, this suggests that tolerance might involve distinct pathways that are more complex to characterize than resistance.

**Table 1.**
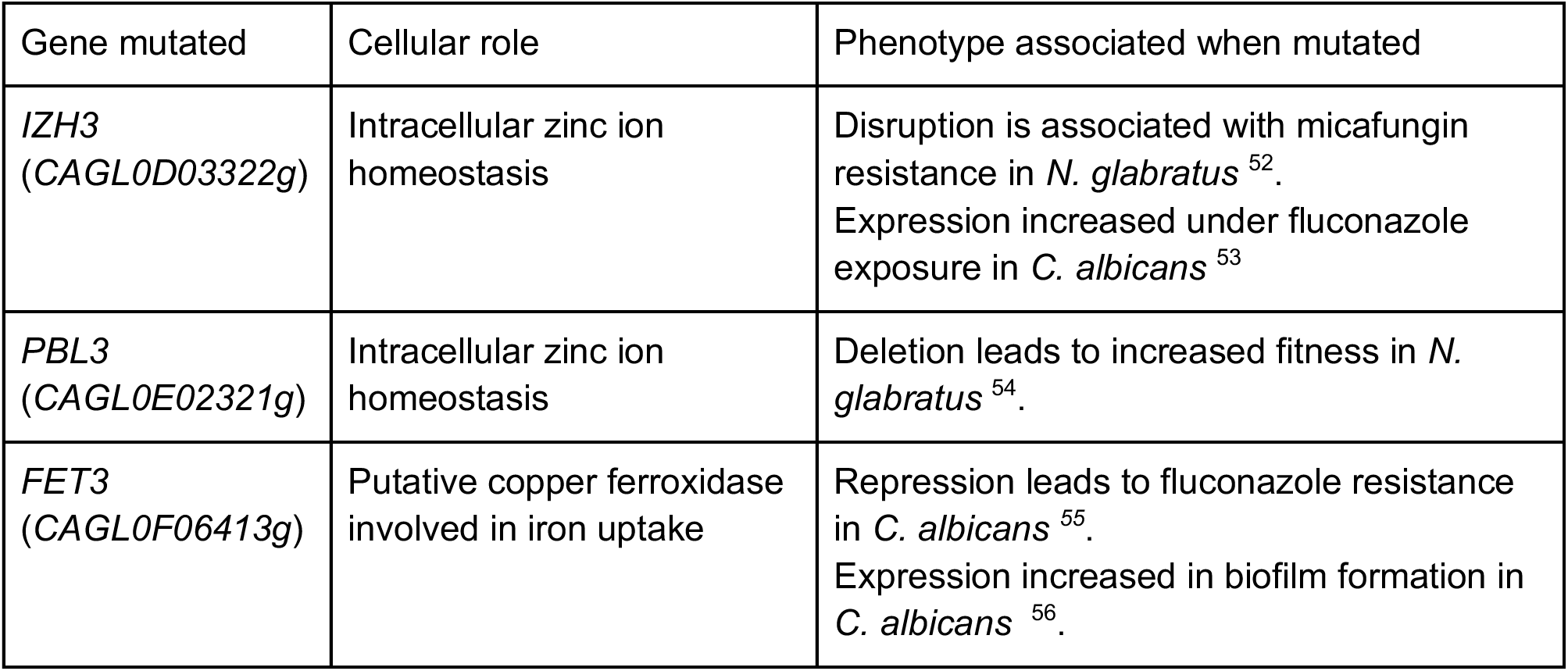
Mutated genes identified in strain WT-FLZ-1-sg and linked to antifungal resistance, increased fitness, or biofilm formation in Candida species.

| Gene mutated | Cellular role | Phenotype associated when mutated |
|---|---|---|
| *IZH3* (*CAGL0D03322g*) | Intracellular zinc ion homeostasis | Disruption is associated with micafungin resistance in *N. glabratus* <sup>52</sup>.<br>Expression increased under fluconazole exposure in *C. albicans* <sup>53</sup> |
| *PBL3* (*CAGL0E02321g*) | Intracellular zinc ion homeostasis | Deletion leads to increased fitness in *N. glabratus* <sup>54</sup>. |
| *FET3* (*CAGL0F06413g*) | Putative copper ferroxidase involved in iron uptake | Repression leads to fluconazole resistance in *C. albicans* <sup>*55*</sup>.<br>Expression increased in biofilm formation in *C. albicans* <sup>56</sup>. |

### Caspofungin adaptation in lig4Δ strains mostly arises through non-canonical genetic pathways

Since CASP resistance is primarily conferred by mutations in the *FKS* genes in *N. glabratus,* particularly at Hot Spot (HS) regions in these genes, we first used targeted sequencing to sequence the two HS regions of both *FKS1* and *FKS2*. Interestingly, WT and *lig4Δ* CASP-evolved strains could be distinguished by the mutations they carry in the HS region of the *FKS* genes (Table 2).

**Table 2.** Targeted-sequencing of *FKS1*, *FKS2*, *ERG3* and *FEN1* of CASP-evolved strains.

| **Strains** | ***FKS1*** |  | ***FKS2*** |  | ***ERG3*** | ***FEN1*** |
|---|---|---|---|---|---|---|
|  | HS1 | HS2 | HS1 | HS2 |  |  |
| WT-CASP-1-sg | - | - | S663F | - | - | - |
| WT-CASP-2-sg | - | - | D666Y | - | - | - |
| WT-CASP-3-sg | - | - | S663F | - | - | - |
| lig4Δ-CASP-1-sg | - | - | - | - | ND | ND |
| lig4Δ-CASP-2-sg | - | - | - | - | ND | ND |
| lig4Δ-CASP-3-sg | - | - | F659del | - | ND | ND |

-: no mutation identified; ND: Not done.

All WT CASP-evolved strains contained one NS mutation in FKS2-HS1, with two out of three strains containing an S663F mutation, which is the most predominant residue mutated in both clinical and *in vitro*-induced echinocandin-resistant *N. glabratus* strains^19,57–59^. We also sequenced CASP-evolved WT strains for *FEN1* and *ERG3,* which are frequently mutated in caspofungin-resistant and fluconazole-cross-resistant strains, respectively^16,60^. However, no mutations in these genes were identified (Table 2).

We identified a mutation in an *FKS* gene (FSK2-F659del in lig4Δ-CASP-3-sg) in only one of the three *lig4Δ*-derived strains (Table 2). Since no mutations in the HS regions of *FKS1* and *FKS2* were identified in the other *lig4Δ* P25-sg strains, we sequenced their whole genomes and examined aneuploidies, SNPs, and INDELs to profile their resistance (Document S2). We also included lig4Δ-CASP-3-sg to further characterize it. No aneuploidy was found in any strain (Fig. S3). However, we identified NS mutations in genes associated with CASP resistance in *N. glabratus*, as summarized in Table 3.

**Table 3.** Summary of mutated genes involved in caspofungin resistance found in *lig4Δ* P25-sg strains.

| Strain | Gene mutated | Cellular role | Phenotype associated when mutated |
|---|---|---|---|
| lig4Δ-CASP-1-sg | *LEM3* (*CAGL0D02442g*) | Putative phospholipase B | Deletion leads to a slight increase in caspofungin resistance in *N. glabratus* <sup>61</sup>. |
| lig4Δ-CASP-2-sg | *SUR4* (*CAGL0G04851g*) | Predicted fatty acid elongase involved in production of very long chain fatty acids for sphingolipid biosynthesis | Deletion confers resistance to caspofungin in *N. glabratus* <sup>61</sup>. |
| lig4Δ-CASP-3-sg | *FKS2* (*CAGL0K04037g*) | 1,3-beta-D-glucan synthase | FSK2-F659del confers caspofungin resistance in *N. glabratus* <sup>62–64</sup>. |
|  | *DFN1* *(CAGL0L11814g)* | Putative glycosylceramide flippase activity, phosphatidylcholine flippase activity, phosphatidylethanolamine flippase activity | Deletion confers caspofungin resistance in *N. glabratus* <sup>65</sup>. |

Cross-adaptation in strains lig4Δ-CASP-1-sg and lig4Δ-CASP-2-sg is likely genetically encoded, as it remains stable after several passages under non-selective conditions (Table S7). Fluconazole cross-adaptation in lig4Δ-CASP-2-sg could be explained by nonsense mutations in *ERG3* and *CAGL0D03102g* (*the S. cerevisiae* ortholog of *COX16*) (Document S2). Cox16 has a putative role in mitochondrial cytochrome c oxidase assembly, and its disruption leads to fluconazole resistance in *N. glabratus* ^51^. *ERG3* encodes a sterol Δ5,6-desaturase, and its mutation or deletion confers fluconazole resistance in *N. glabratus*^16,66–68^. We did not identify mutations in azole-resistance genes in lig4Δ-CASP-1-sg (Document S2). Together, these results demonstrate that caspofungin resistance and fluconazole cross-resistance can arise through multiple genetic routes. Furthermore, they suggest that NHEJ deficiency may constrain access to canonical *FKS*-mediated resistance mechanisms, thereby favouring the emergence of alternative adaptive trajectories.

### lig4Δ drug-evolved populations exhibit a distinct protein signature of drug adaptation

Compared with genomic and transcriptional mechanisms of antifungal resistance, changes in protein abundance during adaptation remain less extensively characterized, although proteomic studies have previously identified some resistance-associated changes in *N. glabratus*^45,49^. We therefore used quantitative proteomics to characterize protein abundance in the P25 drug-evolved populations relative to their non-evolved ancestors. We analyzed whole populations rather than individual P25-sg isolates to capture population-level responses and heterogeneity that may be obscured when analyzing single colonies. All evolved populations exhibited changes in protein abundance, ranging from a single differentially abundant protein in WT-CASP-2 to 81 in lig4Δ-CASP-2, with a median of 24.5 differentially abundant proteins across the 12 populations (Fig. 6A; Document S3). However, the total number of differentially abundant proteins did not differ consistently between FLZ- and CASP-evolved populations or between WT and *lig4Δ* genetic backgrounds (Fig. 6A).

**Figure 6.**
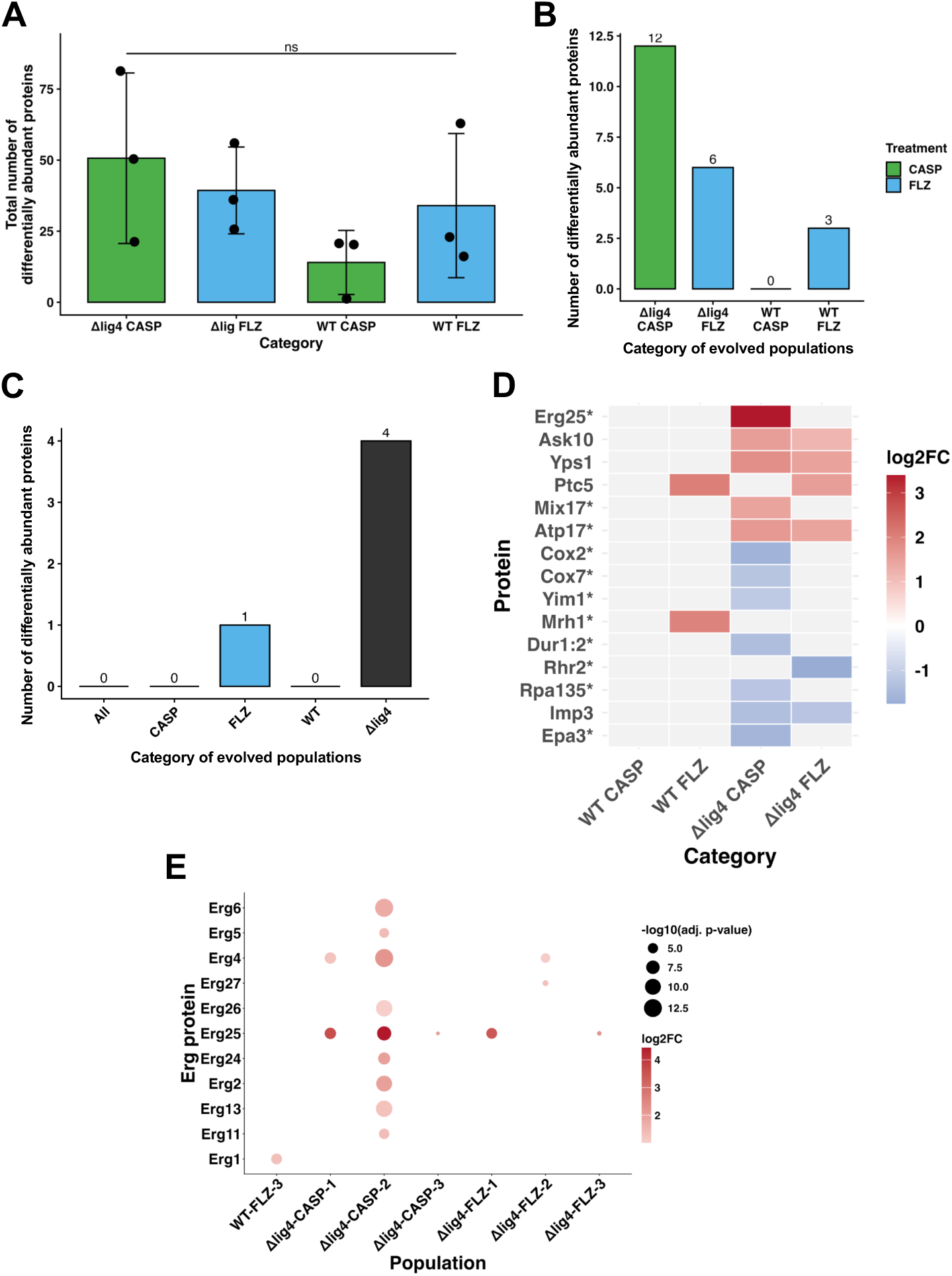
Proteome analysis of drug-evolved P25 populations. **(A)** Total number of differentially abundant proteins in drug-evolved P25 populations. ns : not significant, ordinary one-way ANOVA. **(B, C)** Number of shared differentially abundant proteins per category (same drug evolved in or same genetic background). Only proteins that were significantly regulated in the same direction of all populations of a category (e.g, upregulation in all of CASP-evolved populations). **(D)** Heatmap of proteins that were consistently differentially abundant within the population categories shown in panel B. Asterisks indicate proteins that were also differentially abundant in at least one population outside the category represented. For example, Erg25p was consistently more abundant in all *lig4Δ* CASP-evolved populations but was also found to be more abundant in the *lig4Δ*-FLZ-1 and *lig4Δ*-FLZ-3 populations. **(E)** Upregulation of Erg proteins in drug-evolved P25 populations. Only populations in which at least one Erg protein was significantly upregulated are shown. Dot colour represents the magnitude of protein abundance change, with darker red indicating higher abundance. Dot size represents statistical significance, with larger dots corresponding to lower adjusted *P-*values.

We next asked whether particular proteomic responses were shared across independently evolved populations within each genetic background and drug condition. *lig4Δ* populations shared a larger number of consistently differentially abundant proteins than WT populations (Fig. 6B). Across both antifungal treatments, four proteins were consistently differentially abundant in all six drug-evolved *lig4Δ* populations (Fig. 6C). Three of these (Ask10p, Imp3p, and Yps1p) were consistently increased in abundance and were not differentially abundant in any WT evolved population (Fig. 6D). Yps1p is involved in maintaining cell-wall integrity in *N. glabratus*, and deletion of *YPS1* increases susceptibility to caspofungin and micafungin both *in vitro* and *in vivo*^69^. In addition, CASP-evolved *lig4Δ* populations, but not CASP-evolved WT populations, shared altered abundance of several mitochondrial proteins, including Cox2p, Cox7p, Mix17p, and Yim1p (Fig. 6D). Given the established involvement of mitochondrial function in echinocandin tolerance and resistance in *N. glabratus*^70–72^, these results suggest that mitochondrial remodelling may contribute to caspofungin adaptation when NHEJ is impaired.

Distinct proteomic signatures were also associated with FLZ adaptation and cross-resistance to FLZ. The mitochondrial protein phosphatase Ptc5p was consistently increased in abundance in all FLZ-evolved populations, irrespective of genetic background, but was not differentially abundant in CASP-evolved populations (Fig. 6D). Since in *C. albicans*, deletion of *PTC5* increases susceptibility to fluconazole^73^, this could suggest that increased Ptc5p abundance may contribute to FLZ adaptation in *N. glabratus*. We also observed increased abundance of multiple ergosterol biosynthesis proteins in FLZ-evolved populations, including Erg1p, Erg2p, Erg4p, Erg5p, Erg6p, Erg11p, Erg13p, Erg24p, Erg26p, and Erg27p, (Fig. 6E). Notably, a similar signature was observed in lig4Δ-CASP-1 and lig4Δ-CASP-2, whose corresponding single-colony isolates displayed stable cross-resistance to FLZ. Previous proteomic studies have implicated remodelling of the ergosterol pathway in azole resistance in *N. glabratus* ^49^. Together, these findings suggest that increased abundance of ergosterol biosynthesis proteins may contribute to both direct FLZ adaptation and FLZ cross-resistance following caspofungin evolution.

## Discussion

In this study, we demonstrate that the non-homologous end-joining (NHEJ) DNA repair pathway contributes to rapid and efficient adaptation to caspofungin in *N. glabratus*. To our knowledge, this is the first study implicating NHEJ in antifungal adaptation in this species. Analysis of caspofungin-evolved strains revealed an apparent difference in mutational patterns between WT and *lig4Δ* backgrounds. All WT strains (3/3) carried a nonsynonymous mutation in *FKS2*, whereas only one of the three *lig4Δ* strains harboured a mutation in an *FKS* gene. Given the limited number of evolved strains analyzed per genetic background, we cannot rule out the possibility that this difference arose by chance. Nevertheless, echinocandin resistance in *N. glabratus* is overwhelmingly associated with nonsynonymous mutations within the conserved hotspot regions of *FKS1* and *FKS2*^74^, and this difference was accompanied by reduced fitness and survival of *lig4Δ* populations during caspofungin evolution. Together, these findings are consistent with the hypothesis that NHEJ deficiency constrains access to canonical *FKS*-mediated resistance and promotes alternative, potentially less efficient evolutionary trajectories. A direct role for NHEJ in generating *FKS* mutations appears unlikely, as NHEJ typically generates insertions or deletions, whereas most resistance-associated *FKS* variants are nonsynonymous substitutions. Instead, NHEJ may indirectly facilitate survival under caspofungin stress, increasing the likelihood that highly beneficial *FKS* mutations subsequently arise and become fixed. Thus, NHEJ may influence which evolutionary trajectories are accessible rather than directly generating resistance mutations.

Consistent with this model, *lig4Δ* drug-evolved populations displayed distinct genetic and proteomic signatures. Caspofungin-resistant *lig4Δ* isolates lacking *FKS* hotspot mutations instead carried mutations in other genes previously associated with echinocandin susceptibility, supporting the idea that caspofungin adaptation can involve mechanisms beyond canonical *FKS* mutations^16^. At the population level, *lig4Δ* drug-evolved populations also showed recurrent increases in Yps1p abundance, suggesting that cell-wall maintenance pathways may be particularly important for adaptation in an NHEJ-deficient background. *YPS1* is partially co-regulated by Slt2^75^, whose overexpression has previously been shown to confer caspofungin resistance^76^. CASP-evolved *lig4Δ* populations additionally exhibited altered abundance of several mitochondrial proteins, including Cox2p, Cox7p, Mix17p, and Yim1p. Given the established involvement of mitochondrial function in echinocandin tolerance and resistance^70–72^, these findings suggest that mitochondrial remodelling may also contribute to caspofungin adaptation when NHEJ is impaired. Together, our genomic and proteomic data support a model in which caspofungin resistance can arise through multiple adaptive mechanisms, with the relative frequency of these different evolutionary trajectories influenced by DNA repair capacity.

Another interesting finding was that half of the caspofungin-resistant strains (3/6) exhibited cross-resistance to fluconazole despite never having been exposed to it. Cross-adaptation to fluconazole has previously been reported in *N. glabratus*, including in anidulafungin-adapted strains where it was frequently associated with *ERG3* mutations ^16^. In our study, several ergosterol biosynthesis proteins were consistently more abundant not only in FLZ-evolved populations but also in the cross-resistant lig4Δ-CASP-1 and lig4Δ-CASP-2 populations. One of these isolates also carried a nonsense mutation in *ERG3*, therefore putatively suggesting that remodelling of the ergosterol biosynthesis pathway may contribute to fluconazole cross-resistance following echinocandin adaptation. Importantly, cross-resistance was not always as stable as resistance to the selecting drug, suggesting that secondary drug-response phenotypes can be more transient. We also observed substantial phenotypic heterogeneity within several evolved populations, with susceptible or tolerant cells persisting even in populations classified as resistant after 25 days of continuous drug exposure. This highlights the value of examining whole populations alongside individual isolates, as analysis of only a small number of colonies may underestimate the diversity of adaptive states present during antifungal evolution.

Our longitudinal experimental design also allowed us to capture the rapid dynamics and physiological consequences of antifungal adaptation. Most laboratory evolution studies in *N. glabratus* have focused on resistance phenotypes after relatively longer periods of passaging^13,16,29,30^, whereas we observed fluconazole adaptation within approximately three days and caspofungin adaptation within approximately six days. Notably, most populations ultimately tolerated drug concentrations higher than those used during evolution. Adaptation also carried drug-specific fitness consequences. For example, FLZ-resistant strains exhibited growth defects under several conditions, including reduced growth in glucose-containing media and increased sensitivity to oxidative stress. These findings are consistent with previously reported fitness costs associated with antifungal resistance in *N. glabratus*^16,77^ and with increased oxidative-stress sensitivity associated with fluconazole resistance in other *Candida* species^78^. In contrast, caspofungin-evolved strains generally exhibited fewer growth defects, with WT-CASP-2-sg showing little apparent fitness cost despite its increased caspofungin resistance. These results demonstrate that different adaptive trajectories can carry substantially different physiological costs, which may influence the persistence of resistance when selective pressures change.

Overall, our findings identify NHEJ as a key determinant of the efficiency and evolutionary trajectory of caspofungin adaptation in *N. glabratus*. Loss of *LIG4* reduced the ability of populations to adapt efficiently to caspofungin and was associated with a shift away from the canonical *FKS*-mediated route toward alternative genetic and proteomic responses. At the same time, our results demonstrate that antifungal adaptation can generate heterogeneous populations, cross-resistance to unrelated antifungal classes, and markedly different fitness trade-offs. Further work will be required to determine precisely how NHEJ facilitates efficient caspofungin adaptation and to establish whether the genetic trajectories observed here extend across diverse *N. glabratus* backgrounds. More broadly, understanding the cellular processes that shape which resistance mechanisms are evolutionarily accessible may help identify therapeutic approaches that constrain the emergence of resistance or exploit the fitness costs associated with particular adaptive routes.

## Material and Methods

### Strains, primers and media

*N. glabratus* strains and primers used in this study are listed in **Table S8** and **Table S9**, respectively. The *lig4Δ* strain was kindly provided by the Fairhead laboratory. Complete deletion of *LIG4* was confirmed by whole-genome sequencing. Strains were grown in broth or on plates at 37°C in YPD (non-selective, 1% Yeast Extract, 1% Peptone, 2% Glucose), YPG (respiration selection, 1% Yeast Extract, 1% Peptone, 2% Glycerol), W0+ura (minimal medium, 0.67 % Yeast Nitrogen Base without amino acids with ammonium sulphate, 2 % glucose, supplemented with uracil), or RPMI-1640 supplemented with 3.5 % morpholinepropanesulfonic acid (MOPS) and 2 % glucose.

### Adaptative laboratory experiments (ALEs)

We conducted experimental evolution experiments using a batch serial transfer approach (Fig.1). Initial susceptibility of strains was monitored using minimum inhibitory concentration (MIC) assays (see below). Strains were passaged for 25 days (∼300 strains) in drugs corresponding to twice their MIC50 to study drug adaptation under a high selective pressure. WT and *lig4*Δ strains (Table S8) were collected from glycerol stocks, plated on YPD plates, and let grown for 48 hrs at 37°C to obtain single colonies. For each strain, a single colony from this plate was used to inoculate 4mL of liquid YPD and was incubated overnight, at 37°C with a shaking of 250 rpm. The next day, these overnight cultures were used to inoculate 500 µL of YPD supplementing with the following: i) 640 µL/mL of fluconazole (FLZ) (Sigma-Aldrich, cat# PHR1160), ii) 32 ng/mL of caspofungin (CASP) (Sigma-Aldrich, SML0425-5MG) or iii) no drugs (control), at a final cell concentration of 2.5×10^5^ cells/mL. This was done in triplicate for each condition and strain (18 samples+1 blank) in a 96-deep well plate and strains were organized in a checkerboard style to avoid cross contamination (Fig.1) and the plate was covered by a membrane (Breathe-Easy® sealing membrane) and one layer of aluminum foil, ensuring optimal oxygenation while limiting evaporation. Samples were grown at 37°C with 60 % humidity in a plate shaking incubator with a shaking of 800 rpm for 25 days, with one passage every day. Adaptation to the drugs involved passages of 50µL of the samples to a fresh (450µL) medium (dilution 1/10) every 24 ± 1 hr. For each passage, media with antifungal drugs were freshly prepared right before the passage using a frozen stock of the drugs. Before each passage, part of the culture was frozen and stored at −80 °C (100 µL of the sample in 100 µL of 50 % glycerol) and OD_600nm_ of each sample was read. The passaging concluded after 25 days, each day corresponding to one passage, producing a total of 2 strains x 3 conditions x 3 replicates x 25 days = 450 passaged/evolved strains. After the last passage, we selected, stored and analyzed a single colony that was picked from agar plates and regrown in liquid medium without drugs (final evolved strains called P25-sg for P25 individual colonies). Note here that we use the term passage to describe the number of days that cells spent in the evolution experiment.

To assess the stability of acquired resistance/tolerance in our evolved populations, P25-sg strains evolved in FLZ and CASP were passaged in YPD for 6 days using a similar passaging protocol as the one described above in “2.2 Laboratory micro-evolution experiment”. After the last passage, we stored and analyzed the last passage of all populations for FLZ and CASP susceptibility by running MIC assays.

### Minimum Inhibitory Concentration (MIC) assays

MIC assays were performed in at least three replicates following the EUCAST E.DEF 7.3.2. protocol ^79^, with the following modifications. MICs were run either on a single colony freshly cultured overnight or on a whole population as a patch of cells diluted in sterile water and, in YPD or in RPMI+Ura media (Table 4).

**Table 4.** Types of cells used to run MICs.

| **Strains** | **Type of cells** | **Medium** | **Reason** |
|---|---|---|---|
| Ancestor strains, WT and *lig4*Δ | Single colonies | RPMI+Ura | Check susceptibility of strains before the evolution experiment |
|  |  | YPD | Assess susceptibility of strains in YPD to run the evolution experiment in 2xMIC50 in YPD |
| Evolved strains | Whole population and single colonies | YPD | Assess susceptibility of the evolved strains |

For MICs run on single colonies, fresh YPD cultures made from a single colony were diluted to an OD_530_ of 0.01 (∼2.10^5^ cells) in sterile miliQ water and used as is to run MICs. For MICs run on whole populations, evolved/passaged strains were patched from the glycerol stock on YPD plates and let grown for 48 hrs at 37°C. A portion of the patch was then diluted in sterile miliQ water, adjusted to an OD_530_ of 0.01 (∼2.10^5^ cells) and used as is to run MICs.

In both cases, MIC plates were incubated at 37°C without shaking and absorbance values at 530 nm were read after 24 h using an Infinite 200 PRO microplate reader (Tecan). As the evolution experiment was performed in YPD medium, MIC assays on evolved/passaged strains were performed in YPD. We defined resistant strains as evolved strain with an MIC50 at least twice as high as the MIC50 of the ancestor strain. Depending on strains, culture conditions and passages, the range of concentrations tested were i) in RPMI+Ura: 480-0.46875 ng/mL for anidulafungin and micafungin, 4-0.0039 µg/mL for CASP and, 64-0.0625 µg/mL for FLZ, ii) in YPD: 125-0.1 ng/mL; 500-0.5 ng/mL; 2-0.00195 µg/mL; 4-0.0039 µg/mL; 8-0.0078 µg/mL or 16-0.015 µg/mL for CASP and, 1,280-1.25 µg/mL; 2,560-2.5 µg/mL or 5,120-5 µg/mL for FLZ.

### Growth curves assays

Fitness of passaged/evolved strains were assessed using growth curves assays. Fresh overnight cultures of P25-sg strains were diluted to an OD_600nm_ of 0.5 in a sterile phosphate-buffered saline (PBS) solution and 20µL of this cell dilution were used to inoculate 180µL of different media (YPD, YPG or W0+ura), in at least three replicates, in a 96-well plate, giving rise to a final OD_600nm_ of 0.05. We then topped cultures with 25µL of mineral oil to prevent evaporation and plates were placed, without lid, in an Infinite 200 PRO microplate reader (Tecan) for 24 hrs at 37°C.

### Serial dilution spotting assays

Sensitivity of evolved strains to different stresses were assessed using serial dilution spotting assays. Fresh overnight cultures of P25-sg strains were diluted in 10-fold serial dilutions in sterile PBS solutions and a drop of each dilution was spotted using a using a 96-pin pinning tool onto YPD agar plates or YPD agar plates containing either 10 mM H_2_0_2_, 1 M NaCl, 100 µg/mL Calcofluor White (CW), 3 mM or 10 mM caffeine. Cells were also spotted on YPD agar plate buffered at pH 3, 5 or 10. Plates were then photographed after 48 hrs at 37 °C.

### Biofilm assays

Biofilm assays were performed as previously described ^75^, the only minor modification being that biofilm assays were run in YPD medium. The ‘relative biofilm formation’ was calculated as follows: (OD_490nm_/OD_600nm_ planktonic cells)_condition_/(OD_490nm_/OD_600nm_ planktonic cells)_ancestor strain_ where the condition strain represents the passaged/evolved strain and the ancestor strain its non evolved parent strain, WT or *lig4Δ*.

### C. elegans infection assays

*C. elegans* strain Au37 (glp-4(bn2);sek-1(km4)) was obtained from the *Caenorhabditis* Genetics Center (CGC) and maintained on superfood nematode growth media (NGM) with *E. coli* (OP50) at 15°C, as previously described ^40^. After 5 days, worms were transferred to 25 C for 2 days to synchronize growth. *N. glabratus* strains: *lig4Δ*-FLZ-2, *lig4Δ* non ev, WT-CASP-3, WT non ev, were inoculated in 2 mL of YPD and incubated overnight at 37°C with shaking at 250 rpm. Infection plates of *N. glabratus* were prepared by plating 10 µL of each overnight culture, in triplicate, onto 35-mm brain heart infusion (BHI) agar plates containing 50 µg/mL kanamycin. Infection plates were incubated at 37°C for 16 h. A solid media-based *C. elegans* infection assay protocol was used, adapted from previous studies ^40^. *C. elegans* were washed twice from NGM plates using 6 mL of S basal and transferred to a 15 mL conical tube. The worms were centrifuged at 1500xg for 30s and washed twice more with fresh S basal. Worms were pipetted onto unseeded NGM plates and 45 worms were picked and dropped onto previously prepared infection plates under a stereomicroscope at x10 magnification. Infection plates with worms were incubated at 25 C and scored for survival every 24 hours. Worms were considered dead when they did not respond to being touched with a pick.

### Target-region sequencing

P25-sg strains evolved in CASP were sequenced for the *FKS1* and *FKS2* Hot Spots (HS) regions, and the *FEN1* gene. We also sequenced the *ERG3* gene for P25-sg WT-CASP strains. Strains were streaked on YPD plates and grown for 48 hrs at 37℃ and a single colony was used to run colony PCRs. Regions/genes were amplified using the the Q5® High-Fidelity DNA Polymerase and primers FKS1-F/FKS1-R, FKS2-F/FKS2-R, FEN1-F/FEN1-R, ERG3-1F/ERG3-1R and ERG3-2F/ERG3-2R for *FKS1*, *FKS2*, *FEN1,* the first part of *ERG3* part 1 and the second part of *ERG3*, respectively. PCR products were cleaned-up using the Monarch® PCR & DNA Cleanup Kit and were then Sanger sequenced.

### Whole Genome Sequencing (WGS)

The non-evolved ancestral WT and *lig4Δ* strains, together with their derived drug-evolved P25-sg isolates, including FLZ-evolved strains (WT and *lig4Δ*) and CASP-evolved *lig4Δ* strains, were subjected to whole-genome sequencing (WGS) at SeqCenter in Pittsburgh, PA, USA. Genomic DNA was extracted using the ZymoBIOMICS DNA Extraction Kit. Whole-genome sequencing was performed using an Illumina 650 Mbp sequencing package (approximately 4.33 million reads per sample), and variants were identified using the breseq pipeline configured for haploid genomes ^76^. Mutations were identified by comparing sequence variants present in drug-evolved strains with those detected in their corresponding ancestral strains. Only variants located within coding regions were considered for further analysis.

### Protein extraction and proteomics

Protein extraction was adapted from a protocol in *Klebsiella pneumoniae* ^77^. P25-sg strains were cultured from a single colony overnight in triplicate in YPD at 37 °C. 1 mL of overnight culture each was pelleted at 3500 x RPM for 5 minutes and washed with 1 mL of filter-sterilized 1 X PBS twice. All samples were kept on ice during extractions. To begin, volumes of 100mM Tris HCl (pH 8.5) buffer were prepared with one Complete Ultra proteinase inhibitor cocktail (PIC) tablet per 10 mL, and 300μL of the dissolved solution was added to each dry pellet. To each sample, 30μL of 20 % sodium dodecyl sulfate (SDS) was added. Cells were then lysed with a probe sonicator and transferred to Eppendorf Protein LoBind microcentrifuge tubes. To each sample, a 1:100 volume of 1 M dithiothreitol (DTT) was added, mixed, and incubated at 95°C for 10 minutes with slight shaking. Once cooled, a 1:10 volume of 0.55 M iodoacetamide (IAA) was added to each sample and incubated at room temperature for 20 min in the dark. Proteins were then precipitated in acetone (final concentration of 80 %) at -20 °C overnight, pelleted at 13,500 X RPM for 10 min, and washed with 500 μL of 80 % acetone twice. Pellets were then left to air dry, and resuspended in 200 μL of 8 M urea + 40 mM HEPES. Finally, protein concentrations were quantified with a bovine serum albumin (BSA) tryptophan assay.

### TMT Labelling Methods

#### Peptide Digestion and Tandem Mass Tag (TMT) Labeling

To prepare the peptides for mass spectrometry, 100 mg of each sample was made to 5% Sodium Docecyl Sulfate (SDS, MP Biochemicals, Solon OH) in a total volume of 100ml. The samples were processed using an on-filter digestion protocol using S-Trap mini spin columns (Protifi, Fairport NY USA) as described in the “S-Trap mini spin column digestion protocol” from Protifi. Chemicals used were Tris(2-carboxyethyl)phosphine (TCEP), Methy methanethiosulfonate MMTS), Triethylammonium bicarbonate (TEAB), MgCl2 - Thermo Scientific Rockford IL, Phosphoric acid, and acetonitrile – Fisher Chemical Ottawa ON, Formic acid - Merek, Darmstadt Germany, Benzonase - EMD Millipore/Novagen, St. Louis MO. The samples were digested with Trypsin (Pierce Trypsin, Protease, MS grade Cat # 90058). The digested N. glabratus samples were concentrated to near-dryness (1-5 ml) using a vacuum centrifuge then were labeled with TMT 16 and 18plex Isobaric Label Reagent Set (Thermo Fisher Scientific, Waltham, USA, cat# A44522 and A52045) as per the manufacturer’s protocol, and labelled samples for each TMT experiment were mixed one-to-one.

#### Liquid Chromatography and Mass Spectrometry (LC-MS/MS)

TMT mixed samples were fractionated into 12 fractions by high-pH C18 reversed-phased liquid chromatography using an Agilent 1260 Infinity liquid chromatography system (Agilent Technologies, Santa Clara, CA) equipped with a XBridge C18 3.5mm, 2.1×100mm column (Waters, Wexford Ireland). Each fraction was concentrated to near-dryness using vacuum centrifugation, resuspended in 80 µL of buffer A (2% acetonitrile, 0.1% formic acid).

### Mass Spectrometry

Each sample was separately analysed using a nano-flow Easy nLC-1200 connected in-line to an Orbitrap Fusion Lumos mass spectrometer with a nanoelectrospray ion source at 2.1 kV (Thermo Fisher Scientific, San Jose, CA, USA). The peptide samples were loaded (2 µl) onto a C18-reversed phase Easy Spray column (50 cm long, 75 µm inner diameter, 2 µm particles, (Thermo Fisher Scientific, San Jose, CA, USA) with 100% buffer A (2% acetonitrile, 0.1% formic acid), and then separated on the same column. Peptides were eluted using a linear gradient of 3-27% buffer B (80% acetonitrile, 0.1% formic acid) over 100 min, 28%-38% buffer B for 15 minutes, 38%-100% buffer B for 5 minutes and a wash at 100% B for 10 minutes at a constant flow rate of 200 nl/min. Total LC/MS/MS run-time was about 150 minutes, including the loading, linear gradient, column wash, and the equilibration.

For data acquisition over a 2 second interval we would dynamically choose the most abundant precursor ions from each survey scan, isolate each in the quadrupole (1.2 _m/z_ isolation width) and fragment by CID (35% normalized collision energy) as many as possible in that 1 second. Each fragment ion scan would also have an MS3 scan preformed using 10 SPS precursors and fragmented by HCD (65% normalized collision energy). The survey scans were acquired in the Orbitrap over a range of _m/z_ 375-1500 with a target resolution of 120000 at _m/z_ 200, the subsequent fragment ion scans were acquired in the Iontrap at the Turbo scan rate, and the MS3 scans were acquired in the Orbitrap with a target resolution of 50000 at m/z 200.

### Proteomics Data Analysis

Raw TMT mass spectrometry data was analyzed with the FragPipe (v 23.1) suite of proteomic data analysis tools. The built-in work flow “TMT16-MS3” was used to analyze each TMT experiment separately. The only modification made to this workflow was changing the label type from TMT-16 to TMT-18 when analyzing the 18plex TMT experiment. Data was searched against a Candida glabrata (NCBI taxid: 284593) fasta database downloaded from Uniprot (www.uniprot.org) on 2026-03-04. Briefly, this method uses quantification at the MS3 level with identification from MS2 ion trap fragmentation data. PSMs are validated with Percolator (v 3.7.1) and TMT-Integrator (v 6.1.1) gives the final quantitative results. In total, 2411 proteins were quantified across all samples. The “abundance_protein.tsv” final output files from FragPipe were used for further data analysis in R.

To combine multiple TMT experiments, the ComBat function from the sva function was used for batch correction of protein intensity values between samples before differential protein expression analysis using the limma package. A contrast matrix was first generated using the makeContrasts function. This contrast matrix was used to create a linear model (contrasts.fit) followed by empirical Bayes (eBayes) analysis to compare protein expression between Candida strains. The final limma results were generated with the topTable method. Results were considered significant for adjusted p-value < 0.05 and log-fold-change > 2. For more information about the specific comparisons made and results from limma analysis see Supplemental info.

### Statistics, figures and ploidy analysis

All graphs and statistical analysis were performed using Graphpad Prism v10.0.0 or Rstudio version 2026.05.1+225. Ploidy status of strains was assessed using Ymap analysis tool (http://lovelace.cs.umn.edu/Ymap/) ^78^.

## Acknowledgements

We thank the members of the Shapiro lab for supportive advice and discussion on the project and Pr. Cécile Fairhead for sharing the *lig4Δ* strain. This work was supported by a Natural Sciences and Engineering Research Council of Canada (NSERC) Discovery Grant (RGPIN-2026-04298) and a Canadian Institutes for Health Research Project Grant (PJT206047). RSS holds the Canada Research Chair (Tier II) in Microbial Functional Genomics. LM was supported by an EvoFunPath NSERC Collaborative Research and Training Experience fellowship. NG. and MF are supported by NSERC CGS-D awards.

## Supplementary Figures

**Fig. S1.**
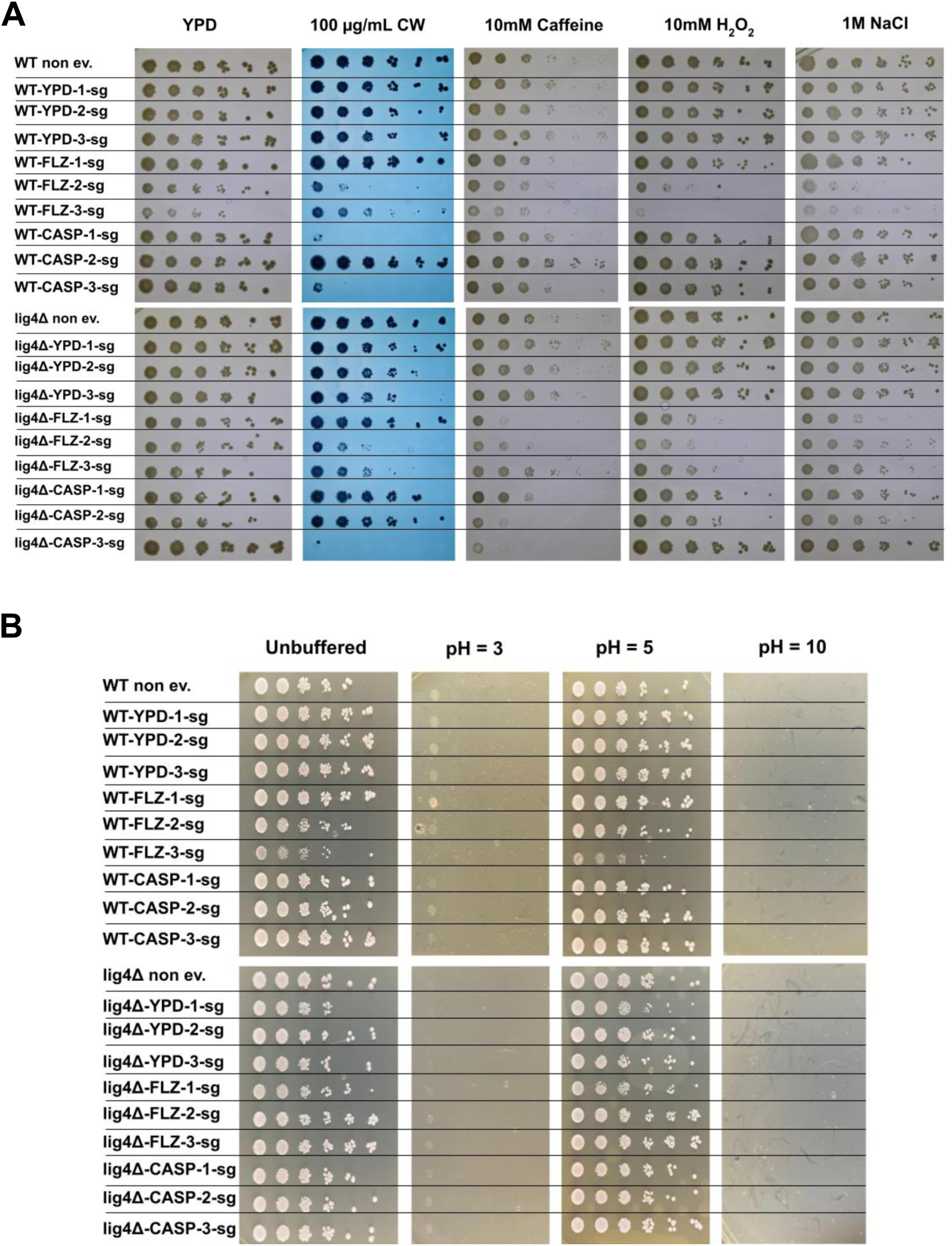
Impact of drug resistance on the stress tolerance of P25-sg strains. Stress tolerance of P25-sg strains was assessed using serial dilution spot assays. Tenfold serial dilutions of cells were spotted onto YPD supplemented with various stressors (A) or adjusted to different pH values (B). The stressors included the cell wall stressors calcofluor white (CW) and caffeine, the oxidative stressor H₂O₂, and the osmotic stressor NaCl. Plates were incubated at 37 °C for 48 h before being photographed.

**Fig. S2.**
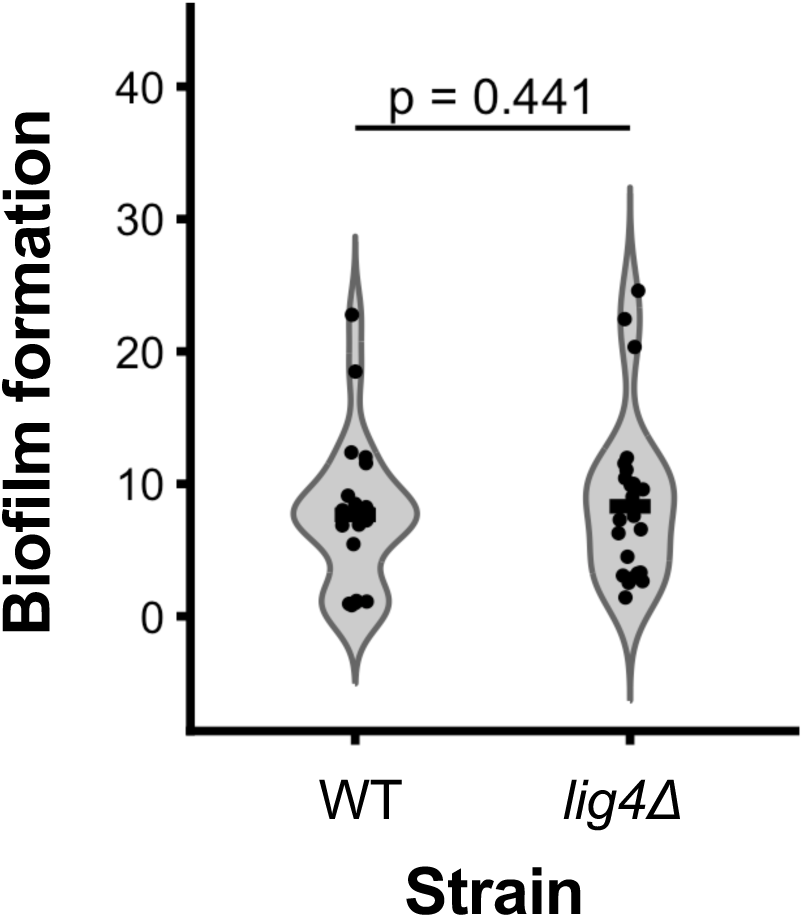
Biofilm formation of the ancestral strains WT and *lig4Δ*. Biofilm formation was assessed using the tetrazolium salt (XTT) reduction assay in YPD. Biofilm formation was calculated as follows: OD_490nm_/OD_600nm_ planktonic cells. Dots represent individual biological replicates from three independent experiments, each containing 8 technical replicates (n=24 total). p = 0.441, Mann-Whitney test.

**Fig. S3.**
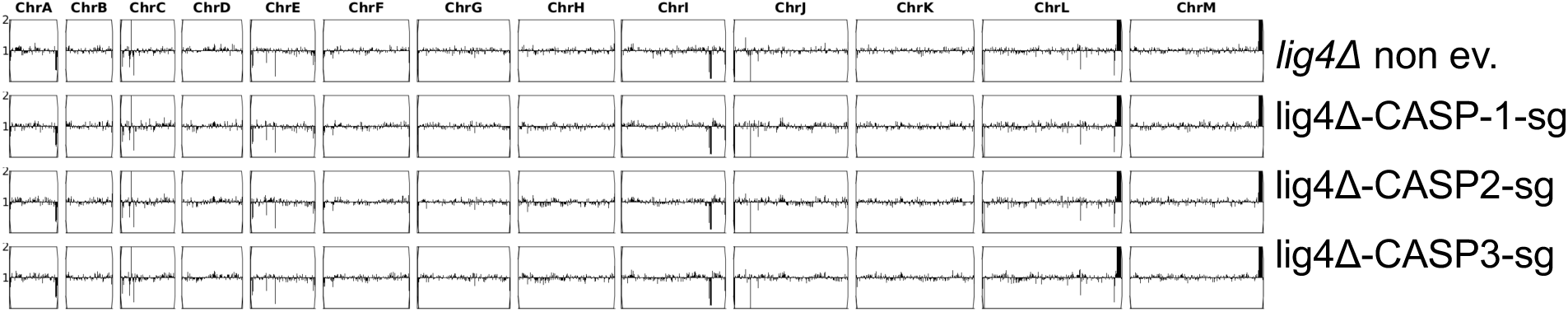
Genomic characterization of drug resistance in P25-sg strains. (A, B) Comparison of the types of mutations (non-synonymous SNPs and INDELs) (A) and the number of mutations per chromosome (B) between drug-evolved strains from the two genetic backgrounds (WT and *lig4Δ*). Statistical analysis in (B) was performed using the Kruskal–Wallis test. (C) Chromosome copy number was determined using the Ymap analysis tool. Black rectangles indicate whole-chromosome duplications.

## Supplementary Tables

**Table S1.**
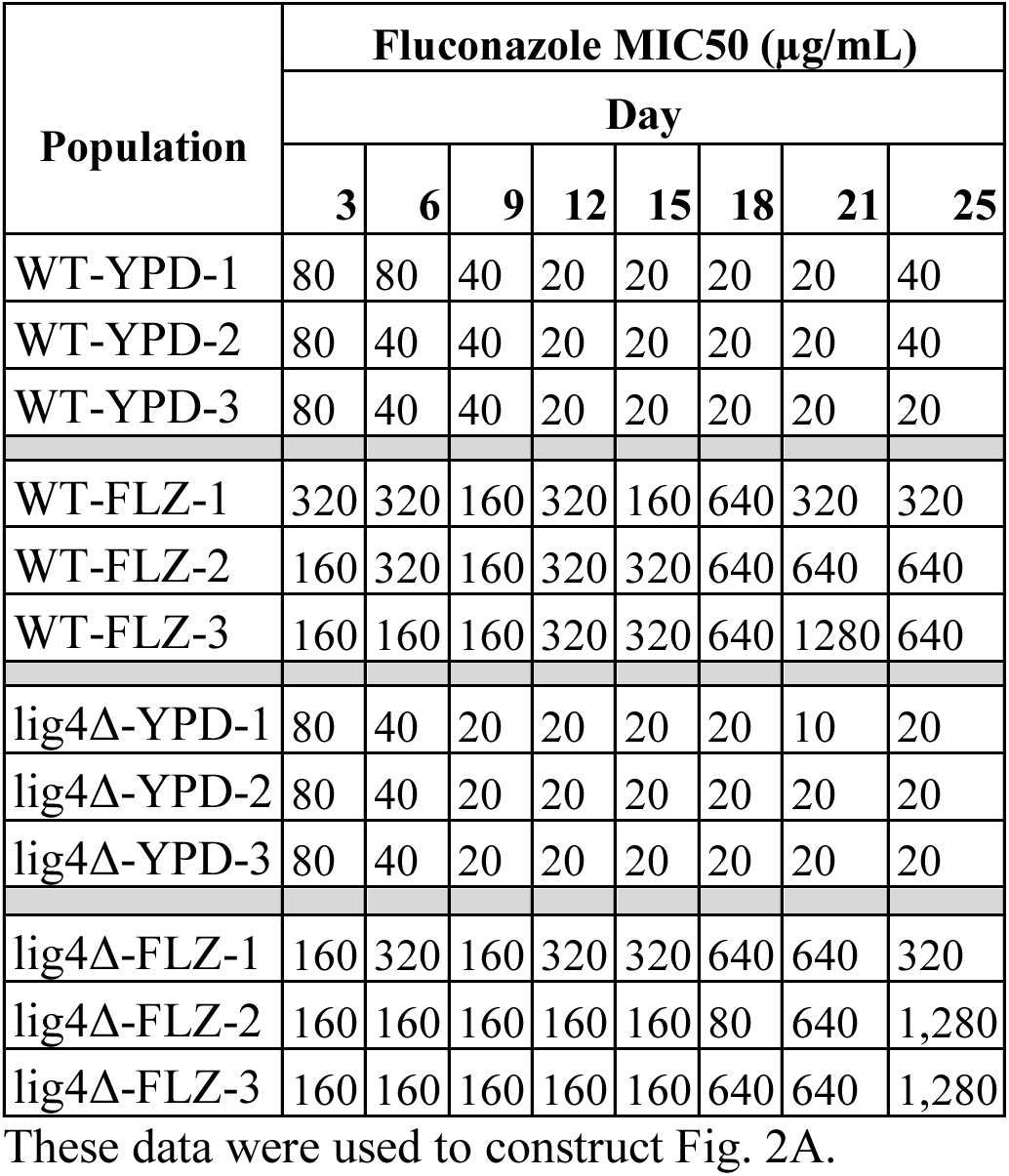
Fluconazole MIC50 of neutrally passaged/evolved populations over time.

**Table S2.** Caspofungin MIC50 of neutrally passaged/evolved populations over time.

| **Population** | **Caspofungin MIC50 (ng/mL)** |  |  |  |  |  |  |  |
|---|---|---|---|---|---|---|---|---|
|  | **Day** |  |  |  |  |  |  |  |
|  | **3** | **6** | **9** | **12** | **15** | **18** | **21** | **25** |
| WT-YPD-1 | 15.625 | 7.8125 | 15.625 | 15.625 | 7.8125 | 7.8125 | 15.625 | 15.625 |
| WT-YPD-2 | 7.8125 | 7.8125 | 7.8125 | 15.625 | 7.8125 | 7.8125 | 7.8125 | 7.8125 |
| WT-YPD-3 | 7.8125 | 7.8125 | 7.8125 | 15.625 | 7.8125 | 7.8125 | 7.8125 | 7.8125 |
| WT-CASP-1 | 7.8125 | 31.25 | 62.5 | 125 | 250 | 250 | 250 | 250 |
| WT-CASP-2 | 7.8125 | 31.25 | 62.5 | 62.5 | 62.5 | 62.5 | 125 | 62.5 |
| WT-CASP-3 | 15.625 | 31.25 | 4,000 | 4,000 | 8,000 | 125 | 125 | 250 |
| lig4Δ-YPD-1 | 7.8125 | 7.8125 | 7.8125 | 15.625 | 7.8125 | 7.8125 | 7.8125 | 7.8125 |
| lig4Δ-YPD-2 | 7.8125 | 7.8125 | 7.8125 | 15.625 | 7.8125 | 7.8125 | 7.8125 | 7.8125 |
| lig4Δ-YPD-3 | 7.8125 | 7.8125 | 7.8125 | 15.625 | 7.8125 | 7.8125 | 7.8125 | 7.8125 |
| lig4Δ-CASP-1 | 15.625 | 16 | 62.5 | 62.5 | 62.5 | 62.5 | 62.5 | 62.5 |
| lig4Δ-CASP-2 | 31.25 | 31.25 | 250 | 250 | 250 | 250 | 250 | 125 |
| lig4Δ-CASP-3 | 15.625 | 16 | 62.5 | 2,000 | 4,000 | 4,000 | 4,000 | 4,000 |

**Table S3.** MIC50 of ancestral strains (non-evolved strains) used to perform the evolution experiment.

| **Strain** | **Medium** | **MIC50 in antifungal drugs** |  |  |  |
|---|---|---|---|---|---|
|  |  | **Fluconazole (µg/mL)** | **Micafungin (ng/mL)** | **Anidulafungin (ng/mL)** | **Caspofungin (ng/mL)** |
| WT | RPMI+Ura | 8 | 30 | 30 | 15.625 |
|  | YPD | 80 | N/D | N/D | 15.625 |
| *lig4Δ* | RPMI+Ura | 8 | 30 | 30 | 15.625 |
|  | YPD | 80 | N/D | N/D | 15.625 |

N/D: Not Done.

**Table S4.** Raw OD600nm values of caspofungin-evolved populations over the first six days of evolution experiments.

| **Population/Day** | **1** | **2** | **3** | **4** | **5** | **6** |
|---|---|---|---|---|---|---|
| **WT-CASP-1** | 0.615100261 | 0.5975000262 | 0.8106999993 | 0.614199996 | 0.6119999886 | 0.689899981 |
| **WT-CASP-2** | 0.6086000204 | 0.6029000282 | 0.731400013 | 0.682099998 | 1 | 0.9595000148 |
| **WT-CASP-3** | 0.364199996 | 0.3849999905 | 0.8457000256 | 0.6880999804 | 0.6335999966 | 0.7074000239 |
| **WT-CASP-4** | 0.6424999833 | 0.6144000292 | 0.5695000291 | 0.6751000285 | 0.7630000114 | 0.7293000221 |
| **WT-CASP-5** | 0.09399999678 | 0.6378999949 | 0.5123000145 | 0.5698999763 | 0.7275999784 | 0.7400000095 |
| **WT-CASP-6** | 0.6678000093 | 0.6104000211 | 0.6281999946 | 0.7225999832 | 0.753099978 | 0.7714999914 |
| **WT-CASP-7** | 0.1000000015 | 0.1932999939 | 0.07689999789 | 0.2910999954 | 0.6516000032 | 0.6750000119 |
| **WT-CASP-8** | 0.6557999849 | 0.613499999 | 0.6308000088 | 0.7027999759 | 0.7833999991 | 0.7558000088 |
| **WT-CASP-9** | 0.09700000286 | 0.6919999719 | 0.1351999938 | 0.6904000044 | 0.7623000145 | 0.7358999848 |
| **WT-CASP-10** | 0.07039999962 | 0.7081999779 | 0.6377000213 | 0.6506000161 | 0.7088999748 | 0.701399982 |
| **WT-CASP-11** | 0.07760000229 | 0.1247000024 | 0.6680999994 | 0.5884000063 | 0.7937999964 | 0.7594000101 |
| **WT-CASP-12** | 0.07010000199 | 0.06759999692 | 0.6887000203 | 0.6108999848 | 0.6385999918 | 0.5916000009 |
| **WT-CASP-13** | 0.07419999689 | 0.08139999956 | 0.6521999836 | 0.6784999967 | 0.8054000139 | 0.7171999812 |
| **WT-CASP-14** | 0.07389999926 | 0.2630999982 | 0.5845000148 | 0.570299983 | 0.6191999912 | 0.6963000298 |
| **WT-CASP-15** | 0.6556000113 | 0.5737000108 | 0.5842999816 | 0.6811000109 | 0.7336000204 | 0.7537000179 |
| **WT-CASP-16** | 0.07630000263 | 0.1004000008 | 0.6164000034 | 0.6101999879 | 0.685199976 | 0.7322999835 |
| **WT-CASP-17** | 0.6777999997 | 0.6058999896 | 0.5989000201 | 0.681400001 | 0.7516000271 | 0.7332999706 |
| **WT-CASP-18** | 0.07649999857 | 0.7336000204 | 0.6794000268 | 0.6528000236 | 0.7490000129 | 0.71450001 |
| **WT-CASP-19** | 0.6039000154 | 0.5950000286 | 0.5992000103 | 0.7347999811 | 0.8151000142 | 0.757799983 |
| **WT-CASP-20** | 0.07429999858 | 0.06599999964 | 0.3635999858 | 0.829400003 | 0.6340000033 | 0.6319000125 |
| **lig4Δ-CASP-1** | 0.3483999968 | 0.3594999909 | 0.6735000014 | 0.5800999999 | 0.6886000037 | 0.7267000079 |
| **lig4Δ-CASP-2** | 0.07339999825 | 0.5827000141 | 0.7713999748 | 0.6233000159 | 0.5942999721 | 0.6906999946 |
| **lig4Δ-CASP-3** | 0.2208999991 | 0.5526999831 | 0.711499989 | 0.5723999739 | 0.7936999798 | 0.7700999975 |
| **lig4Δ-CASP-4** | 0.6492000222 | 0.5792999864 | 0.6563000083 | 0.7110999823 | 0.7470999956 | 0.7421000004 |
| **lig4Δ-CASP-5\*** | 0.07119999826 | 0.06499999762 | 0.06580000371 | 0.06230000034 | 0.06440000236 | 0.06260000169 |
| **lig4Δ-CASP-6** | 0.6667000055 | 0.5831000209 | 0.6340000033 | 0.706700027 | 0.7767000198 | 0.780099988 |
| **lig4Δ-CASP-7** | 0.07020000368 | 0.06440000236 | 0.06340000033 | 0.1268000007 | 0.4806999862 | 0.4433999956 |
| **lig4Δ-CASP-8** | 0.628000021 | 0.6187999845 | 0.6464999914 | 0.7217000127 | 0.7663999796 | 0.7628999949 |
| **lig4Δ-CASP-9** | 0.07249999791 | 0.07880000025 | 0.638800025 | 0.6128000021 | 0.6927000284 | 0.7182999849 |
| **lig4Δ-CASP-10** | 0.07410000265 | 0.7480999827 | 0.684400022 | 0.7139000297 | 0.7436000109 | 0.7210999727 |
| **lig4Δ-CASP-11** | 0.07280000299 | 0.06599999964 | 0.2180999964 | 0.5821999907 | 0.2013999969 | 0.256099999 |
| **lig4Δ-CASP-12** | 0.07320000231 | 0.7382000089 | 0.6148999929 | 0.6561999917 | 0.6833000183 | 0.6754000187 |
| **lig4Δ-CASP-13** | 0.07050000131 | 0.067900002 | 0.08240000159 | 0.6934999824 | 0.8112000227 | 0.6854000092 |
| **lig4Δ-CASP-14** | 0.07320000231 | 0.06660000235 | 0.6258000135 | 0.7293999791 | 0.6787999868 | 0.7542999983 |
| **lig4Δ-CASP-15\*** | 0.06859999895 | 0.06530000269 | 0.1784999967 | 0.07599999756 | 0.06490000337 | 0.08990000188 |
| **lig4Δ-CASP-16** | 0.07329999655 | 0.7056999803 | 0.6301000118 | 0.6220999956 | 0.7081999779 | 0.7864000201 |
| **lig4Δ-CASP-17** | 0.06989999861 | 0.08169999719 | 0.5881000161 | 0.5961999893 | 0.5870000124 | 0.6168000102 |
| **lig4Δ-CASP-18\*** | 0.06759999692 | 0.06469999999 | 0.06480000168 | 0.06340000033 | 0.06319999695 | 0.06190000102 |
| **lig4Δ-CASP-19\*** | 0.06499999762 | 0.06639999896 | 0.06480000168 | 0.06310000271 | 0.06570000201 | 0.06350000203 |
| **lig4Δ-CASP-20** | 0.07320000231 | 0.7057999969 | 0.6237999797 | 0.5393000245 | 0.6672999859 | 0.6642000079 |
| **Blank for pop1-3** | 0.06710000336 | 0.06570000201 | 0.0671999976 | 0.06560000032 | 0.06549999863 | 0.06610000134 |
| **Blank for pop4-20** | 0.06849999726 | 0.06580000371 | 0.06570000201 | 0.06430000067 | 0.06440000236 | 0.06379999965 |

Shades of red indicate population growth, with darker shades representing higher growth. Populations 4–20 for both the WT and *lig4Δ* strains were evolved at a different time from populations 1–3, but following the same experimental protocol. Blank cells correspond to YPD containing 32 ng/mL fluconazole in both cases. CASP: caspofungin. \*: Populations that died during the evolution experiment.

**Table S5.** MIC50 values of single colonies from FLZ-evolved P25 populations (P25-sg)

| **Population** | **FLZ MIC50 (µg/mL)** |  |  |  |  |  |  |  |  |  |  |
|---|---|---|---|---|---|---|---|---|---|---|---|
|  | **P25 population** | **Single colonies from P25 (P25-sg)** |  |  |  |  |  |  |  |  |  |
|  |  | **#1** | **#2** | **#3** | **#4** | **#5** | **#6** | **#7** | **#8** | **#9** | **#10** |
| **WT-FLZ-1** | 320 | 40 | 40 | 40 | 40 | N/A | N/A | N/A | N/A | N/A | N/A |
| **WT-FLZ-2** | 640 | 320 | N/A | N/A | N/A | N/A | N/A | N/A | N/A | N/A | N/A |
| **WT-FLZ-3** | 640 | 320 | N/A | N/A | N/A | N/A | N/A | N/A | N/A | N/A | N/A |
| ***lig4*Δ–FLZ-1** | 320 | 20 | 10 | 10 | 10 | 20 | 20 | 10 | 10 | 10 | 10 |
| ***lig4*Δ–FLZ-2** | 1,280 | 640 | N/A | N/A | N/A | N/A | N/A | N/A | N/A | N/A | N/A |
| ***lig4*Δ–FLZ-3** | 1,280 | 80 | 640 | 640 | 640 | 640 | N/A | N/A | N/A | N/A | N/A |

MIC50 values of P25 populations were already presented in Table S3; they were included in this table for a clearer comparison between the MIC50 of P25 populations and single colonies derived from these populations (P25-sg). Strains with an MIC50>80µg/mL are considered resistant. For all populations, colony #1 (colony #2 for *lig4*Δ–FLZ-3) were chosen to follow up with. FLZ: Fluconazole, P25: Population 25, N/A: Not applicable.

**Table S6.** MIC50 values of single colonies from CASP-evolved P25 populations (P25-sg)

|  | **CASP MIC50 (ng/mL)** |  |  |  |  |  |  |  |  |  |  |
|---|---|---|---|---|---|---|---|---|---|---|---|
| **Population** | **P25 population** | **Single colonies from P25 (P25-sg)** |  |  |  |  |  |  |  |  |  |
|  |  | **#1** | **#2** | **#3** | **#4** | **#5** | **#6** | **#7** | **#8** | **#9** | **#10** |
| **WT-CASP-1** | 250 | 62.5 | 62.5 | 31.25 | 7.8125 | 31.25 | 31.25 | 15.625 | 31.25 | 125 | 31.25 |
| **WT-CASP-2** | 62.5 | 31.25 | N/A | N/A | N/A | N/A | N/A | N/A | N/A | N/A | N/A |
| **WT-CASP-3** | 250 | 125 | N/A | N/A | N/A | N/A | N/A | N/A | N/A | N/A | N/A |
| **lig4Δ–CASP-1** | 62.5 | 62.5 | N/A | N/A | N/A | N/A | N/A | N/A | N/A | N/A | N/A |
| **lig4Δ–CASP-2** | 125 | 15.625 | 250 | 31.25 | N/A | N/A | N/A | N/A | N/A | N/A | N/A |
| **lig4Δ–CASP-3** | 4,000 | 4,000 | N/A | N/A | N/A | N/A | N/A | N/A | N/A | N/A | N/A |

MIC50 values of P25 populations were already presented in Table S4 and are included here to facilitate comparison with the MIC50 values of P25 populations and single colonies derived from these populations (P25-sg). Strains with an MIC50>15.625 ng/mL are considered resistant. Colonies picked to follow up with were: colony #1 for WT-CASP-2, WT-CASP-3, *lig4*Δ–CASP-1, *lig4*Δ–CASP-3; colony #9 for WT-CASP-1 and colony #2 for *lig4*Δ–CASP-2. CASP: Caspofungin, P25: Population 25, N/A: Not applicable.

**Table S7.** Resistance and cross-resistance of P25-sg and P25-sg-YPD strains to fluconazole and caspofungin.

| Population strains are from | Fluconazole MIC50 (µg/mL) |  | Caspofungin MIC50 (ng/mL) |  |
|---|---|---|---|---|
|  | P25-sg | P25-sg-YPD | P25-sg | P25-sg-YPD |
| WT-FLZ-1 | 40 | 40 | 7.8125 | 15.625 |
| WT-FLZ-2 | 320 (R) | 320 (R) | 7.8125 | 15.625 |
| WT-FLZ-3 | 320 (R) | 640 (R) | 15.625 | 15.625 |
| *lig4*Δ-FLZ-1 | 20 | 20 | 15.625 | 15.625 |
| *lig4*Δ-FLZ-2 | 640 (R) | 640 (R) | 15.625 | 15.625 |
| *lig4*Δ-FLZ-3 | 640 (R) | 640 (R) | 7.8125 | 15.625 |
| WT-CASP-1 | 160 (R) | 20 | 125 (R) | 250 (R) |
| WT-CASP-2 | 80 | 80 | 31.25 (R) | 31.25 (R) |
| WT-CASP-3 | 40 | 10 | 125 (R) | 125 (R) |
| *lig4*Δ-CASP-1 | 1,280 (R) | 2,560 (R) | 62.5 (R) | 62.5 (R) |
| *lig4*Δ-CASP-2 | 1,280 (R) | 640 (R) | 250 (R) | 250 (R) |
| *lig4*Δ-CASP-3 | 20 | 10 | 4,000 (R) | 8,000 (R) |

Fluconazole and caspofungin MIC50 of P25-sg strains were already presented in Table S7 and S8 and are included here to facilitate comparison with the MIC50 values of P25-sg and P25-sg-YPD strains. P25-sg-YPD: P25-sg strains passaged in YPD for six days. FLZ: fluconazole, CASP: caspofungin. (R): Resistance.

**Table S8.** Strains used in this work.

| **Strains** | **Parent** | **Genotype** | **Reference** |
|---|---|---|---|
| CBS138 | N/A | *MATalpha LIG4* | PMID: 15229592 |
| HM100 | CBS138 | *MATalpha ura3Δ::KANMX LIG4* | PMID: 18375614 |
| YL02 | HM100 | *MATalpha ura3Δ::KANMX LIG4* | PMID: 33057400 |
| YL07 WT (WT strain) | YL02 | *MATalpha-inc ura3Δ::KANMX LIG4* | PMID: 33057400 |
| YL07 *lig4*Δ (*lig4*Δ strain) | YL02 | *MATalpha-inc ura3Δ::KANMX lig4*Δ | Kind donation from the Fairhead lab |

N/A: Not Applicable.

**Table S9.**
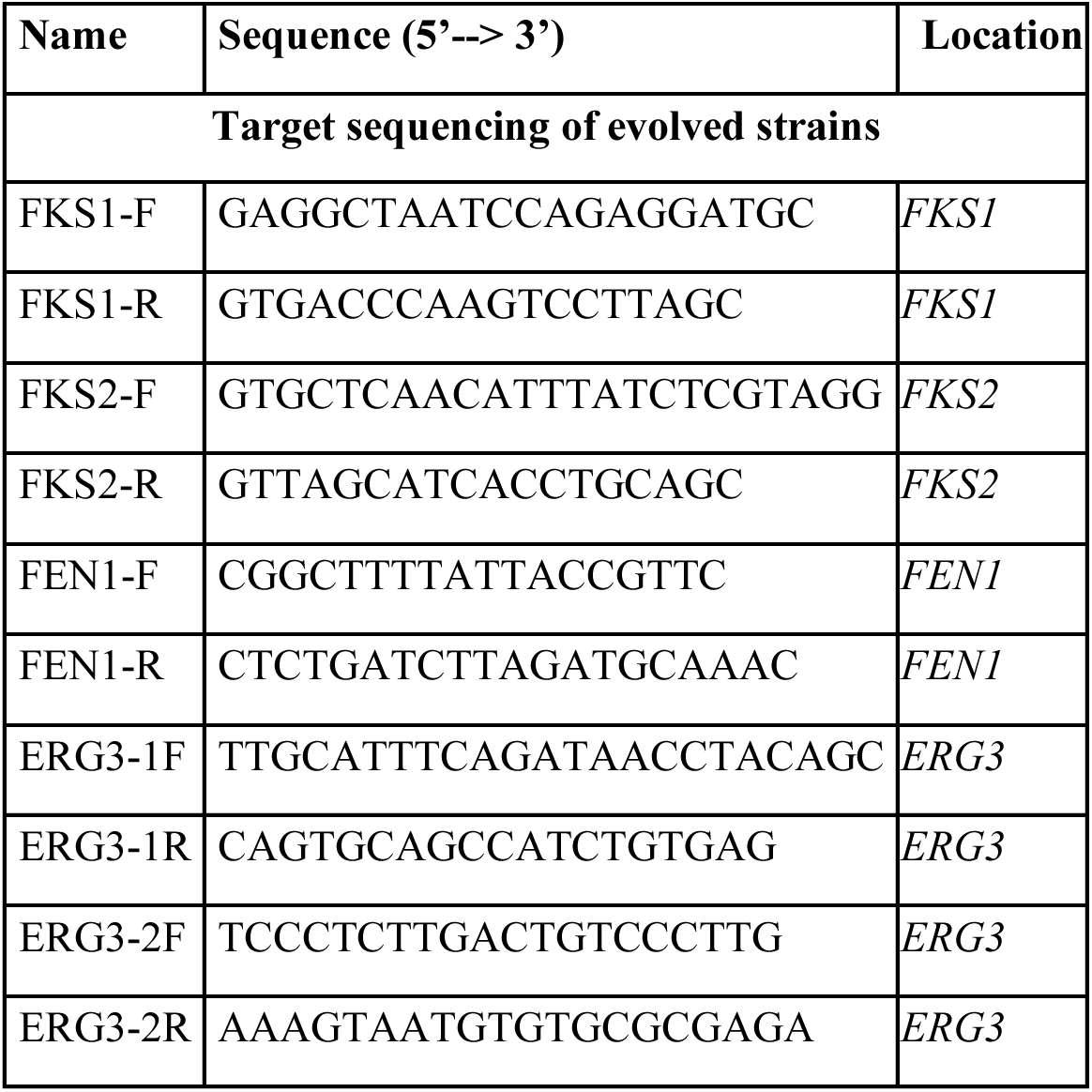
Primers used in this work.

## Notes

### Competing Interest Statement

The authors have declared no competing interest.

## References

1. Denning, D. W. Global incidence and mortality of severe fungal disease. Lancet Infect. Dis. 24, e428–e438 (2024).

2. Oliveira, L. V. N., Wang, R., Specht, C. A. & Levitz, S. M. Vaccines for human fungal diseases: close but still a long way to go. NPJ Vaccines 6, 33 (2021).

3. Hoenigl, M. et al. The Antifungal Pipeline: Fosmanogepix, Ibrexafungerp, Olorofim, Opelconazole, and Rezafungin. Drugs 81, 1703–1729 (2021).

4. Hui, S. T., Gifford, H. & Rhodes, J. Emerging Antifungal Resistance in Fungal Pathogens. Curr Clin Microbiol Rep 11, 43–50 (2024).

5. Vanzolini, T. & Magnani, M. Old and new strategies in therapy and diagnosis against fungal infections. Appl. Microbiol. Biotechnol. 108, 147 (2024).

6. Puumala, E., Fallah, S., Robbins, N. & Cowen, L. E. Advancements and challenges in antifungal therapeutic development. Clin. Microbiol. Rev. 37, e0014223 (2024).

7. Mota Fernandes, C., et al. The Future of Antifungal Drug Therapy: Novel Compounds and Targets. Antimicrob. Agents Chemother. 65, (2021).

8. Beardsley, J. et al. Candida glabrata (Nakaseomyces glabrata): A systematic review of clinical and microbiological data from 2011 to 2021 to inform the World Health Organization Fungal Priority Pathogens List. Med. Mycol. 62, (2024).

9. Duggan, S. & Usher, J. Candida glabrata: A powerhouse of resistance. PLoS Pathog. 19, e1011651 (2023).

10. Shields, R. K., Nguyen, M. H., Press, E. G., Updike, C. L. & Clancy, C. J. Caspofungin MICs correlate with treatment outcomes among patients with Candida glabrata invasive candidiasis and prior echinocandin exposure. Antimicrob. Agents Chemother. 57, 3528– 3535 (2013).

11. Whaley, S. G. et al. Azole Antifungal Resistance in Candida albicans and Emerging Non-albicans Candida Species. Front. Microbiol. 7, 2173 (2016).

12. Salazar, S. B., et al. Disclosing azole resistance mechanisms in resistant Candida glabrata strains encoding wild-type or gain-of-function CgPDR1 alleles through comparative genomics and transcriptomics. G3 12, (2022).

13. Galocha, M. et al. Genomic evolution towards azole resistance in Candida glabrata clinical isolates unveils the importance of CgHxt4/6/7 in azole accumulation. Commun Biol 5, 1118 (2022).

14. Vu, B. G., Stamnes, M. A., Li, Y., Rogers, P. D. & Moye-Rowley, W. S. The Candida glabrata Upc2A transcription factor is a global regulator of antifungal drug resistance pathways. PLoS Genet. 17, e1009582 (2021).

15. Poláková, S. et al. Formation of new chromosomes as a virulence mechanism in yeast *Candida glabrata*. Proc. Natl. Acad. Sci. U. S. A. 106, 2688–2693 (2009).

16. Ksiezopolska, E. et al. Narrow mutational signatures drive acquisition of multidrug resistance in the fungal pathogen Candida glabrata. Curr. Biol. 31, 5314–5326.e10 (2021).

17. Alexander, B. D. et al. Increasing echinocandin resistance in Candida glabrata: clinical failure correlates with presence of FKS mutations and elevated minimum inhibitory concentrations. Clin. Infect. Dis. 56, 1724–1732 (2013).

18. Pham, C. D. et al. Role of FKS Mutations in Candida glabrata: MIC values, echinocandin resistance, and multidrug resistance. Antimicrob. Agents Chemother. 58, 4690–4696 (2014).

19. Misas, E. et al. Genomic description of acquired fluconazole- and echinocandin-resistance in patients with serial Candida glabrata isolates. J. Clin. Microbiol. 62, e0114023 (2024).

20. Naskar, S., Prajapati, A. & Kaur, R. Antifungal drug resistance in Candida glabrata: role of cellular signaling and gene regulatory networks. FEMS Yeast Res. 25, (2025).

21. Legrand, M., Chan, C. L., Jauert, P. A. & Kirkpatrick, D. T. Role of DNA mismatch repair and double-strand break repair in genome stability and antifungal drug resistance in Candida albicans. Eukaryot. Cell 6, 2194–2205 (2007).

22. Healey, K. R. et al. Prevalent mutator genotype identified in fungal pathogen Candida glabrata promotes multi-drug resistance. Nat. Commun. 7, 11128 (2016).

23. Agyare-Tabbi, M. R. et al. The putative error prone polymerase REV1 mediates DNA damage and drug resistance in Candida albicans. NPJ Antimicrob. Resist. 2, 42 (2024).

24. Emerson, C. H. & Bertuch, A. A. Consider the workhorse: Nonhomologous end-joining in budding yeast. Biochem. Cell Biol. 94, 396–406 (2016).

25. Maroc, L. & Fairhead, C. A new inducible CRISPR-Cas9 system useful for genome editing and study of double-strand break repair in Candida glabrata. Yeast 36, 723–731 (2019).

26. Cen, Y., Fiori, A. & Van Dijck, P. Deletion of the DNA ligase IV gene in Candida glabrata significantly increases gene-targeting efficiency. Eukaryot. Cell 14, 783–791 (2015).

27. Carreté, L. et al. Patterns of genomic variation in the opportunistic pathogen candida glabrata suggest the existence of mating and a secondary association with humans. Curr. Biol. 32, 3219 (2022).

28. Belenky, P., Camacho, D. & Collins, J. J. Fungicidal drugs induce a common oxidative-damage cellular death pathway. Cell Rep. 3, 350–358 (2013).

29. Ksiezopolska, E., Schikora-Tamarit, M. À., Carlos Nunez-Rodriguez, J. & Gabaldón, T. Long-term stability of acquired drug resistance and resistance associated mutations in the fungal pathogen Nakaseomyces glabratus (Candida glabrata). Front. Cell. Infect. Microbiol. 14, 1416509 (2024).

30. Cavalheiro, M. et al. A Transcriptomics Approach To Unveiling the Mechanisms of In Vitro Evolution towards Fluconazole Resistance of a Candida glabrata Clinical Isolate. Antimicrob. Agents Chemother. 63, (2019).

31. Brauner, A., Fridman, O., Gefen, O. & Balaban, N. Q. Distinguishing between resistance, tolerance and persistence to antibiotic treatment. Nat. Rev. Microbiol. 14, 320–330 (2016).

32. Shantal, C.-J. N., Juan, C.-C., Lizbeth, B.-U. S., Carlos, H.-G. J. & Estela, G.-P. B. Candida glabrata is a successful pathogen: An artist manipulating the immune response. Microbiol. Res. 260, 127038 (2022).

33. Vincent, B. M., Lancaster, A. K., Scherz-Shouval, R., Whitesell, L. & Lindquist, S. Fitness trade-offs restrict the evolution of resistance to amphotericin B. PLoS Biol. 11, e1001692 (2013).

34. Hill, J. A., O’Meara, T. R. & Cowen, L. E. Fitness trade-offs associated with the evolution of resistance to antifungal drug combinations. Cell Rep. 10, 809–819 (2015).

35. Carolus, H. et al. Acquired amphotericin B resistance leads to fitness trade-offs that can be mitigated by compensatory evolution in Candida auris. Nat. Microbiol. 9, 3304–3320 (2024).

36. Rodrigues, C. F., Rodrigues, M. E., Silva, S. & Henriques, M. Candida glabrata biofilms: How far have we come? J. Fungi (Basel) 3, (2017).

37. Cavalheiro, M. & Teixeira, M. C. Candida biofilms: Threats, challenges, and promising strategies. Front. Med. (Lausanne) 5, 28 (2018).

38. Gupta, P., Gupta, H. & Poluri, K. M. Geraniol eradicates Candida glabrata biofilm by targeting multiple cellular pathways. Appl. Microbiol. Biotechnol. 105, 5589–5605 (2021).

39. Rodríguez-Cerdeira, C. et al. Biofilms and vulvovaginal candidiasis. Colloids Surf. B Biointerfaces 174, 110–125 (2019).

40. Kim, G. H., Rosiana, S., Kirienko, N. V. & Shapiro, R. S. A Simple Nematode Infection Model for Studying Candida albicans Pathogenesis. Current Protocols in Microbiology vol. 59 Preprint at 10.1002/cpmc.114 (2020).

41. Rosiana, S. et al. Comprehensive genetic analysis of adhesin proteins and their role in virulence of Candida albicans. Genetics 10.1093/genetics/iyab003 (2021) doi:10.1093/genetics/iyab003.

42. Pukkila-Worley, R., Peleg, A. Y., Tampakakis, E. & Mylonakis, E. Candida albicans hyphal formation and virulence assessed using a Caenorhabditis elegans infection model. Eukaryot. Cell 8, 1750–1758 (2009).

43. Noble, S. M., French, S., Kohn, L. A., Chen, V. & Johnson, A. D. Systematic screens of a Candida albicans homozygous deletion library decouple morphogenetic switching and pathogenicity. Nat. Genet. 42, 590–598 (2010).

44. vanden Bossche, H., Marichal, P., Odds, F. C., Le Jeune, L. & Coene, M. C. Characterization of an azole-resistant Candida glabrata isolate. Antimicrob. Agents Chemother. 36, 2602–2610 (1992).

45. Bing, J. et al. Experimental Evolution Identifies Adaptive Aneuploidy as a Mechanism of Fluconazole Resistance in Candida auris. Antimicrob. Agents Chemother. 65, (2020).

46. Selmecki, A., Gerami-Nejad, M., Paulson, C., Forche, A. & Berman, J. An isochromosome confers drug resistance in vivo by amplification of two genes, ERG11 and TAC1. Mol. Microbiol. 68, 624–641 (2008).

47. 47. Gervais, N. C. et al. Chromosome-scale CRISPR screening reveals secretory pathway genes as drivers of aneuploidy-mediated antifungal tolerance. bioRxiv 2026.07.23.740334 (2026) doi:10.64898/2026.07.23.740334.

48. Vu, B. G. & Moye-Rowley, W. S. Azole-resistant alleles of ERG11 in Candida glabrata trigger activation of the Pdr1 and Upc2A transcription factors. Antimicrob. Agents Chemother. 66, e0209821 (2022).

49. Rogers, P. D., Vermitsky, J.-P., Edlind, T. D. & Hilliard, G. M. Proteomic analysis of experimentally induced azole resistance in Candida glabrata. J. Antimicrob. Chemother. 58, 434–438 (2006).

50. 50. Vermitsky, J.-P. et al. Pdr1 regulates multidrug resistance in Candida glabrata: gene disruption and genome-wide expression studies. Molecular Microbiology vol. 61 704–722 Preprint at 10.1111/j.1365-2958.2006.05235.x (2006).

51. Gale, A. N. et al. Identification of Essential Genes and Fluconazole Susceptibility Genes in Candida glabrata by Profiling Hermes Transposon Insertions. G3 10, 3859–3870 (2020).

52. Nickels, T. J., Gale, A. N., Harrington, A. A., Timp, W. & Cunningham, K. W. Transposon-sequencing (Tn-seq) of the Candida glabrata reference strain CBS138 reveals epigenetic plasticity, structural variation, and intrinsic mechanisms of resistance to micafungin. G3 14, (2024).

53. Copping, V. M. S. et al. Exposure of Candida albicans to antifungal agents affects expression of SAP2 and SAP9 secreted proteinase genes. J. Antimicrob. Chemother. 55, 645–654 (2005).

54. Schwarzmüller, T. et al. Systematic phenotyping of a large-scale Candida glabrata deletion collection reveals novel antifungal tolerance genes. PLoS Pathog. 10, e1004211 (2014).

55. Rogers, P. D. & Barker, K. S. Evaluation of differential gene expression in fluconazole-susceptible and -resistant isolates of Candida albicans by cDNA microarray analysis. Antimicrob. Agents Chemother. 46, 3412–3417 (2002).

56. Nett, J. E., Lepak, A. J., Marchillo, K. & Andes, D. R. Time course global gene expression analysis of an in vivo Candida biofilm. J. Infect. Dis. 200, 307–313 (2009).

57. Bordallo-Cardona, M. Á. et al. In vitro exposure to increasing micafungin concentrations easily promotes echinocandin resistance in Candida glabrata isolates. Antimicrob. Agents Chemother. 61, AAC.01542–16 (2017).

58. Bordallo-Cardona, M. Á. et al. Low and constant micafungin concentrations may be sufficient to lead to resistance mutations in FKS2 gene of Candida glabrata. Med. Mycol. 56, 903–906 (2018).

59. Aldejohann, A. M., Herz, M., Martin, R., Walther, G. & Kurzai, O. Emergence of resistant Candida glabrata in Germany. JAC Antimicrob. Resist. 3, dlab122 (2021).

60. Hassoun, Y. et al. Evolutionary dynamics in gut-colonizing Candida glabrata during caspofungin therapy: Emergence of clinically important mutations in sphingolipid biosynthesis. PLoS Pathog. 20, e1012521 (2024).

61. Healey, K. R., Katiyar, S. K., Raj, S. & Edlind, T. D. CRS-MIS in Candida glabrata: sphingolipids modulate echinocandin-Fks interaction: Sphingolipids modulate echinocandin-Fks interaction. Mol. Microbiol. 86, 303–313 (2012).

62. Szymankiewicz, M. et al. Case report: Echinocandin-resistance Candida glabrata FKS mutants from patient following radical cystoprostatectomy due to muscle-invasive bladder cancer. Front. Oncol. 11, 794235 (2021).

63. Garcia-Effron, G., Lee, S., Park, S., Cleary, J. D. & Perlin, D. S. Effect of Candida glabrata FKS1 and FKS2 mutations on echinocandin sensitivity and kinetics of 1,3-beta-D-glucan synthase: implication for the existing susceptibility breakpoint. Antimicrob. Agents Chemother. 53, 3690–3699 (2009).

64. Sig, A. K. et al. The emergence of echinocandin-resistant Candida glabrata exhibiting high MICs and related FKS mutations in Turkey. J. Fungi (Basel) 7, 691 (2021).

65. Rosenwald, A. G. et al. Identification of genes in candida glabrata conferring altered responses to caspofungin, a cell wall synthesis inhibitor. G3 (Bethesda) 6, 2893–2907 (2016).

66. Yoo, J. I., Choi, C. W., Lee, K. M. & Lee, Y. S. Gene expression and identification related to fluconazole resistance of Candida glabrata strains. Osong Public Health Res. Perspect. 1, 36–41 (2010).

67. Gregor, J. B., et al. An expanded toolkit of drug resistance cassettes for Candida glabrata, Candida auris, and Candida albicans leads to new insights into the ergosterol pathway. mSphere 8, e0031123 (2023).

68. Pais, P., Galocha, M., Takahashi-Nakaguchi, A., Chibana, H. & Teixeira, M. C. Multiple genome analysis of Candida glabrata clinical isolates renders new insights into genetic diversity and drug resistance determinants. Microb. Cell Fact. 9, 174–189 (2022).

69. Garcia-Rubio, R. et al. Critical assessment of cell wall integrity factors contributing to in vivo echinocandin tolerance and resistance in Candida glabrata. Front. Microbiol. 12, 702779 (2021).

70. Xue, P. et al. Mitochondrial characteristics of echinocandin tolerance and resistance in Candida glabrata and Candida krusei. Front. Microbiol. 17, 1906168 (2026).

71. Garcia-Rubio, R. et al. Multifactorial role of mitochondria in echinocandin tolerance revealed by transcriptome analysis of drug-tolerant cells. MBio 12, e0195921 (2021).

72. Arastehfar, A. et al. Overlooked Candida glabrata petites are echinocandin tolerant, induce host inflammatory responses, and display poor in vivo fitness. MBio 14, e0118023 (2023).

73. Zhao, Y., Feng, J., Li, J. & Jiang, L. Mitochondrial type 2C protein phosphatases CaPtc5p, CaPtc6p, and CaPtc7p play vital roles in cellular responses to antifungal drugs and cadmium in Candida albicans. FEMS Yeast Res. 12, 897–906 (2012).

74. Bédard, C. et al. FungAMR: a comprehensive database for investigating fungal mutations associated with antimicrobial resistance. Nat. Microbiol. 10, 2338–2352 (2025).

75. Maroc, L., Shaker, H. & Shapiro, R. S. Functional genetic characterization of stress tolerance and biofilm formation in Nakaseomyces (Candida) glabrata via a novel CRISPR activation system. mSphere e0076123 (2024).

76. Deatherage, D. E. & Barrick, J. E. Identification of mutations in laboratory-evolved microbes from next-generation sequencing data using breseq. Methods Mol. Biol. 1151, 165–188 (2014).

77. Reitzel, C., Sukumaran, A., Zanetti, C., Muselius, B. & Geddes-McAlister, J. Phosphoproteome Profiling of Klebsiella pneumoniae under Zinc-Limited and Zinc-Replete Conditions. Microbiol Resour Announc 12, e0025823 (2023).

78. Abbey, D. A. et al. YMAP: a pipeline for visualization of copy number variation and loss of heterozygosity in eukaryotic pathogens. Genome Med. 6, 100 (2014).

79. 79. Arendrup, M. C., Meletiadis, J., Mouton, J. W., Lagrou, K. & Hamal, P. EUCAST DEFINITIVE DOCUMENT E.DEF 7.3.2 Method for the determination of broth dilution minimum inhibitory concentrations of antifungal agents for yeasts. https://www.eucast.org/fileadmin/src/media/PDFs/EUCAST_files/AFST/Files/EUCAST_E_Def_7.3.2_Yeast_testing_definitive_revised_2020.pdf.

